# Rampant Reticulation in a Rapid Radiation of Tropical Trees - Insights from *Inga* (Fabaceae)

**DOI:** 10.1101/2023.09.12.557345

**Authors:** Rowan J. Schley, Rosalía Piñeiro, James A. Nicholls, Flávia Fonseca Pezzini, Catherine Kidner, Audrey Farbos, Jens J. Ringelberg, Alex D. Twyford, Kyle G. Dexter, R. Toby Pennington

## Abstract

Evolutionary radiations underlie much of the species diversity of life on Earth, particularly within the world’s most species-rich tree flora – that of the Amazon rainforest. Hybridisation occurs in many radiations, with effects ranging from homogenisation of species to generation of genetic and phenotypic novelty that fuels speciation, but the influence of hybridisation on Amazonian tree radiations has been little studied. We address this using the ubiquitous, species-rich, neotropical tree genus *Inga*, which typifies rapid radiations of rainforest trees. We assess patterns of gene tree incongruence to ascertain whether hybridisation was associated with rapid radiation in *Inga.* Given the importance of insect herbivory in structuring rainforest tree communities (and hence the potential for hybridisation to promote adaptation through admixture of defence traits), we also test whether introgression of loci underlying chemical defences against herbivory occurred during the radiation of *Inga.* Our phylogenomic analyses of 189/288 *Inga* species using >1300 target capture loci showed widespread introgression in *Inga*. Specifically, we found widespread phylogenetic incongruence explained by introgression, with phylogenetic networks recovering multiple introgression events across *Inga* and up to 20% of shared, likely introgressed, genetic variation between some species. In addition, most defence chemistry loci showed evidence of positive selection and marginally higher levels of introgression. Overall, our results suggest that introgression has occurred widely over the course of *Inga’s* history, likely facilitated by extensive dispersal across Amazonia, and that in some cases introgression of chemical defence loci may influence adaptation in *Inga*.

---

Rapid evolutionary radiations that generate exceptionally species-rich groups are a fundamental component of biodiversity (Hughes et al. 2015). Hybridisation (interbreeding between species) is frequent in rapid evolutionary radiations (Seehausen 2004) but its evolutionary role has been long debated. While hybridisation can result in ‘speciation reversal’ that reduces diversity (Vonlanthen et al. 2012; Kearns et al. 2018), or may be ‘lineage-neutral’ and have no effect on diversification (Justison et al. 2023), it is frequently invoked as a catalyst of rapid radiation (e.g. Barrier et al. 1999; Meier et al. 2017; Lamichhaney et al. 2018). This is because hybridisation can ‘reshuffle’ existing genetic variation, generating genomic and phenotypic novelty (Rieseberg et al. 2007; Marques et al. 2019) that may confer adaptation to new environments (e.g. adaptive introgression and/or transgressive segregation (Rieseberg et al. 1999; Suarez-Gonzalez et al. 2018)) or lead to re-sorting of intrinsic incompatibilities (e.g. Bateson-Dobzhansky-Muller incompatibilities) that promote reproductive isolation and hence rapid speciation (Schumer et al. 2015).

It is possible to detect genetic admixture and infer past reticulation events in a clade through examining gene tree conflict (Doyle 1992; Naciri and Linder 2015). The proportions of different conflicting gene tree topologies can indicate the relative contributions of introgression (transfer of genetic material following persistent hybridisation) and incomplete lineage sorting (ILS) to phylogenetic incongruence (e.g. Green et al. 2010; Durand et al. 2011; Pease et al. 2018). This incongruence can also help estimate the relative genetic contributions of progenitor lineages to introgressant descendants (Patterson et al. 2012). There is a growing body of work that explores incongruence to better understand introgression across the tree of life, particularly in plants (e.g. in oaks (McVay et al. 2017) and willows (Wagner et al. 2020)), but only recently have such studies been undertaken in the most species-rich flora on Earth - that of neotropical rainforests (Schley et al. 2020; Larson et al. 2021; reviewed in Schley et al. 2022).

The flora of neotropical rainforests is remarkable in its species diversity (Antonelli and Sanmartín 2011; Ulloa Ulloa et al. 2017; Raven et al. 2020), particularly at local scales – there are more tree species in a single hectare of the Amazon (c. 655 spp. (Valencia et al. 2004)) than in all of Europe (c. 454 spp. (Rivers et al. 2019)). Many species-rich neotropical plant groups arose through recent, rapid radiations (e.g. Erkens et al. 2007, Annonaceae; Koenen et al. 2015, Meliaceae) but the influence of hybridisation on plant radiations has been little studied, and virtually not at all in tropical rainforest trees (reviewed in Abbott (2017)). The prevailing view, based largely on morphological patterns, has been that hybrids between rainforest tree species are exceptionally rare (e.g. Ashton 1969), but this is challenged by recent genomic data for a few Amazonian tree species (e.g. *Brownea* (Schley et al. 2020); *Eschweilera* (Larson et al. 2021)).

The tree genus *Inga* Mill. (Fabaceae) is widespread and abundant in neotropical rainforests, and was the first documented example of rapid radiation in the Amazonian tree flora (Richardson et al. 2001). *Inga* exhibits the highest diversification rate of any Amazonian tree genus (Baker et al. 2014), with *ca.* 300 species arising in the last *ca.* 10 Ma (Ringelberg et al. 2023). Similar recent, rapid radiation events in other tree genera gave rise to a large portion of the Amazonian tree flora - over half of Amazonian tree species belong to genera with >100 species (Dexter and Chave 2016). *Inga* is an ideal study system to understand the influence of hybridisation on rapid rainforest tree radiations due to the large volume of previous work examining the diversification, ecology and biogeography of the group (Richardson et al. 2001; Kursar et al. 2009; Dexter et al. 2017; Forrister et al. 2019) coupled with its ubiquity and species diversity in the Amazon (Pennington, T. D. 1997). Previous phylogenetic work on *Inga,* based on Sanger sequencing of relatively few species, revealed low resolution of species-level relationships (Richardson et al. 2001; Kursar et al. 2009; Dexter et al. 2010), with resolution improving in later phylogenomic studies using 22 *Inga* species (Nicholls et al. 2015). Here we generate the most comprehensive phylogenetic tree of *Inga* to date, comprising >1300 loci and 189 species, greatly improving resolution of *Inga* species relationships to help understand whether hybridisation influenced diversification.

Hybridisation may be more widespread than initially thought in rainforest tree radiations like *Inga*, first and foremost because of their remarkable level of co-occurrence in local rainforest communities. Up to 19 species of *Inga* can coexist in 1ha, and such high local diversity is typical of many other species-rich Amazonian tree genera (e.g. *Protium* and *Eschweilera* (Valencia et al. 1994; Larson et al. 2021)), some of which have emerging evidence of hybridisation (e.g. between 3 *Eschweilera* species in Manaus, Brazil). Furthermore, there is substantial overlap in flowering times of many *Inga* species, which share a wide range of pollinators due to their generalist pollination syndrome (Koptur 1983), and recent work using microsatellites suggested hybridisation occurs between two *Inga* species in Peru (Rollo et al. 2016).

These dispersal-assembled local communities of *Inga* (Dexter et al. 2017) are largely structured by insect herbivore pressure, such that co-occurring *Inga* species differ more in their chemical defences against herbivores than expected by chance (Kursar et al. 2009; Endara et al. 2022). This is because higher densities of conspecifics with the same defences leads to increased mortality from herbivores that can overcome these defences (Janzen 1970; Connell 1971; Forrister et al. 2019). Divergent adaptation in chemical defence traits is documented in *Inga* over evolutionary timescales (Forrister et al. 2023), likely driven by these negative density-dependent processes over ecological timescales (Forrister et al. 2019). This suggests that it is adaptive to possess rare defence chemistry phenotypes, as fewer local herbivores can overcome them. Therefore, the transfer of defence chemistry loci between *Inga* species via hybridisation, followed by positive selection on those loci, may be predicted to facilitate colonisation of new ecological space and eventually lead to speciation. To our knowledge this has never been investigated from a genomic perspective. Therefore, here we aim to:

1. Infer the diversification history of *Inga* by generating the most comprehensive phylogenetic tree of the genus to date;
2. Assess patterns of phylogenetic incongruence and reticulate evolution, resulting from hybridisation, across *Inga’*s evolutionary history;
3. Assess whether hybridisation may have contributed to *Inga*’s rapid diversification through transfer of adaptive genomic loci underlying chemical defence against herbivores.

## Materials & Methods

### Taxon Sampling and DNA Sequencing

We performed phylogenomic analyses on target capture sequencing data from 189 of the 282 accepted *Inga* species (67%), 109 of which were sequenced for this study, and 80 of which were taken from previous work (one from Nicholls et al. 2015; 79 from Ringelberg et al. 2023). Our sampling was based on a list of all accepted *Inga* species compiled using the World Checklist of Vascular Plants (WCVP 2020) (as of June 2021) and a monograph of *Inga* (Pennington, T. D. 1997). Preliminary analyses identified 13 subclades within *Inga*, with the size of these used to ensure proportional per-subclade down-sampling for computationally-intensive downstream analyses.

To contextualise our *Inga* analyses at broader phylogenetic scales, we collated sequence data from many outgroup species generated by previous studies (Koenen et al. 2020; Ringelberg et al. 2022). We included all seven genera and 32 other species from the ‘Inga clade’ (*sensu* Koenen et al. 2020) within which *Inga* is nested, including its sister genus *Zygia*. In addition, we included a further 73 species comprising all 42 genera from the ‘Ingoid clade’ (sensu Koenen et al. 2020) within which the Inga clade is nested. Finally, we included 22 species from the broader Mimosoid legume clade, giving a total of 127 outgroup species. A list of accessions sampled including species, sampling location, data source and voucher information is detailed in Supplementary Table S1, available on Dryad.

DNA was extracted from 20 mg of dried leaf material with the DNeasy Plant Mini Kit (Qiagen, Hilden, Germany) using modifications described in Nicholls et al. (2015). DNA library preparation, enrichment and sequencing were carried out either by Arbor BioSciences (Ann Arbor, MI, USA) or the University of Exeter sequencing service (Exeter, UK) following the NEBnext Ultra II FS protocol (New England Biolabs, Ipswich, MA, USA). Targeted bait capture was performed with a subfamily-specific ‘Mimobaits’ bait set (Nicholls et al. 2015; Koenen et al. 2020) using the MyBaits protocol v.2 and 3 (Arbor Biosciences, Ann Arbor, MI, USA). The Mimobaits set targets 1320 loci, including 113 genes coding for enzymes underlying anti-herbivore defence chemistry in *Inga* (hereafter ‘defence chemistry’ loci). Other loci targeted by the Mimobaits set are ‘single-copy phylogenetically informative’ loci that were selected due to their high levels of informative substitutions and the fact they were single-copy (1044), ‘differentially expressed’ loci that had different numbers of transcriptome reads between the species used to design the baits (109) and ‘miscellaneous’ loci (54), which are unannotated but contain phylogenetic signal. Enriched libraries were sequenced using the NovaSeq 6000 platform with a paired-end 150bp run on two S1 flow cell lanes.

### Sequence Assembly, Trimming and Alignment

All analyses were conducted on the UK Crop Diversity Bioinformatics HPC Resource. DNA sequencing reads were quality-checked with FASTQC 0.11.3 (Andrews 2010) and were trimmed using TRIMMOMATIC 0.3.6 (Bolger et al. 2014) to remove adapter sequences and to quality-filter reads. TRIMMOMATIC settings permitted < 2 mismatches, a palindrome clip threshold of 30 and a simple clip threshold of 10. Bases with a quality score < 28 and reads shorter than 36 bases long were removed from the dataset. Following quality-filtering, reads were mapped to target loci using BWA 0.7.17 (Li and Durbin 2009), loci were assembled using SPADES 3.11.1 (Bankevich et al. 2012) with a coverage cut-off of 5× and exons were extracted with EXONERATE (Slater and Birney 2005), all of which are implemented in the HYBPIPER pipeline 1.2 (Johnson et al. 2016). Recovery of loci was visualised using the ‘gene_recovery_heatmap.R’ script distributed with HYBPIPER. To improve the signal:noise ratio and remove possible paralogues from our *Inga* dataset we used the Putative Paralogs Detection pipeline 1.0.1, (PPD; https://github.com/Bean061/putative_paralog; Zhou et al. (2022)), described in Supplementary Methods on Dryad.

We conducted all subsequent analyses on three data subsets to include a broad range of phylogenetic scales and to assess the influence of paralogy on our analyses. All datasets comprised a single accession per species, the first of which included 189 *Inga* species with *Zygia* sp. ‘*mediana*’ as the outgroup (dataset ‘Singlesp’). The second dataset included 127 outgroup species and proportional per-subclade sampling of 50 *Inga* species (dataset ‘Outgroup’), chosen to include the *Inga* species with the best locus recovery per subclade. The third and final dataset comprised the same 189 accessions as the ‘Singlesp’ *Inga* dataset, but loci were assembled and cleaned using the PPD pipeline, with paralogous loci removed (dataset ‘PPD’).

Targeted loci for each of the three datasets were aligned by gene region using 1000 iterations in MAFFT 7.453 (Katoh and Standley 2013) with the *‘—adjustdirectionaccurately*’ flag to incorporate reversed sequences. These alignments were then cleaned using the ‘-*automated1*’ flag in trimAl 1.3 (Capella-Gutiérrez et al. 2009), and realigned with MAFFT using the ‘*—auto’* flag. This resulted in 1305 refined alignments for the ’Singlesp’ dataset, 1044 for the ‘Outgroup’ dataset and 1267 loci for the ‘PPD’ dataset. Alignment summaries detailing proportions of variable sites and missing data were then generated with AMAS (Borowiec 2016).

### Phylogenomic Analyses

Gene trees were inferred for each locus alignment across the three datasets using IQ-TREE (Nguyen et al. 2015) by selecting the best-fit substitution model (*-MFP*) while reducing the impact of severe model violations (*-bnni*) with 1000 ultrafast bootstrap replicates. Following this, a ‘species tree’ was generated based on the best-scoring IQtrees using ASTRALMP 5.15.5 under the default parameters (Zhang et al. 2018) for all three datasets. Following concatenation of all locus alignments within the ‘Singlesp’ data subset using AMAS, we used the same parameters as above to infer a phylogenetic tree from the concatenated supermatrix using IQ-TREE. Finally, we visualised shared genetic variation among species by building neighbour net plots with uncorrected P-distances in SPLITSTREE v4.14.6 (Huson and Bryant 2005) for each of the three datasets.

### Analysing Incongruence and Reticulation

To assess incongruence among our gene trees, we estimated three metrics implemented in the QUARTET SAMPLING method 1.3.1 (https://www.github.com/fephyfofum/quartetsampling; Pease et al. (2018)) based on each dataset’s ASTRAL species tree, using 100 replicate runs. For each node, we estimated Quartet Concordance (QC) to assess whether there was incongruence, Quartet Differential (QD) to assess whether one incongruent topology was favoured and Quartet Informativeness (QI) to assess whether data were sufficiently informative to distinguish well-supported incongruence from lack of signal (QI not shown as all datasets were informative).

Having assessed incongruence across our three datasets, we then visually investigated gene tree conflict of certain higher-level relationships highlighted in the Quartet Sampling analysis with DISCOVISTA (Sayyari et al. 2018) (subclades defined in Supplementary Table S2, available on Dryad). We interpreted gene tree conflict as “low” if the proportion of gene trees supporting the species tree topology was >30 percent higher than for both alternative, and nodes with one conflicting topology above the 33% ‘equal frequency’ threshold were examined more closely as possibly suggesting introgression after Kuhnhäuser et al. (2021).

We then assessed whether the incongruence we inferred was caused by introgression or ILS using the *Dtrios* function in DSUITE (Malinsky et al. 2021). We used Patterson’s D-statistic (i.e. the ‘ABBA-BABA’ test; Green et al. 2010; Durand et al. 2011) and estimated the proportion of shared variation between species using the F_4_ ratio (Patterson et al. 2012), where for both metrics values closer to 1 indicate more introgression. We generated an input VCF file for *Dtrios* for each of the three datasets by calling SNPs using BWA 0.7.17, SAMTOOLS 1.13 (Danecek et al. 2021) and BCFTOOLS 1.13 (Li 2011) as in Joana Meier’s ‘Speciation Genomics’ github (https://speciationgenomics.github.io), using our target bait set sequences as the reference. The resulting VCFs were filtered to contain sites with > 8x coverage and a quality score >20, as well as removing taxa with >50% total missing data.

For each taxon trio test set, we additionally used the *‘--abbaclustering*’ tool in DSUITE to account for variation in substitution rate across clades, and so more accurately infer introgression without false positives resulting from homoplasy (Koppetsch et al. 2023). To further minimise the effect of substitution rate variation for the Outgroup dataset, we only included Ingoid clade species in our DSUITE analysis, along with genera from its close sister group (*Jupunba/Hydrochorea/Punjuba/Pseudalbizia*), using *Cedrelinga/Pseudosamanea/Chloroleucon/Samanea/Boliviadendron/Enterolobium/Albizia* as outgroup. For the *Inga* Singlesp and PPD analyses, we used the closely-related *Zygia sp. ‘mediana’* as outgroup. We assessed significance of each test using 20 block jackknife resampling runs (ca. 50,000-65,000 variants per block), following which we corrected *D* and ABBAclustering *P-*values for multiple testing with the Benjamini–Hochberg correction in RSTATIX (Kassambara, 2020) using R v 4.2.1 (R Development Core Team 2013). Finally, we filtered out test sets without significant ABBA clustering (indicating homoplasy), and visualized our D-statistic and F_4_ ratio estimates with Ruby scripts available from https://github.com/mmatschiner. D-statistics are best at detecting recent introgression (Bjorner et al. 2022), and older hybridisation events can result in correlated F4 ratios between related species. To infer deeper introgression events we estimated the F_branch_ statistic (Malinsky et al. 2018) for each combination of taxa, filtered results to only include trios with significant ABBAclustering, and plotted scores with the DSUITE ‘dtools.py’ utility.

To model historical reticulation across *Inga* and outgroup species we inferred phylogenetic networks with SNAQ!, implemented in the JULIA v1.7.2 (Bezanson et al. 2017) package PHYLONETWORKS v0.16.2 (Solís-Lemus, Bastide, & Ané, 2017). We inferred phylogenetic networks from representative down-sampled subsets of our ‘Singlesp’ and ‘Outgroup’ datasets with between 0-4 reticulation events (*h*), due to computational limitations, excluding the PPD dataset due to the minimal effect of paralogs on our analyses. We estimated networks by calculating quartet concordance factors (CF) for each node, which were also used to estimate γ values (probabilities of ancestral contribution to hybridization events). The best-fit network was chosen using negative log-pseudolikelihood comparison, selecting the *h-*value above which likelihood scores did not improve *(hmax)*. We performed the same analysis on representative subsets of *Inga* subclades to better understand within-subclade reticulation. Downsampled datasets prioritised accessions with the highest locus recovery, aiming for proportional sampling of subclades. Details of D and F statistics, as well as accession selection for *PHYLONETWORKS,* are found in Supplementary Methods on Dryad.

### Assessing Per-Locus Introgression and Selection

We assessed whether defence chemistry loci experienced elevated introgression and selection relative to other loci in *Inga*. To do this we first estimated the per-locus proportion of introgression using the *f _dM_* statistic (Malinsky et al. 2015) in DSUITE. *f _dM_* more accurately infers introgression in small genomic windows than *D-*statistics (Martin et al. 2014; Malinsky et al. 2015), while using the same sampling design (i.e., three taxa and an outgroup). We performed *f _dM_* analysis on three ‘subclade subsets’, which were selected based on inferred introgression events from PHYLONETWORKS and DSUITE. Each analysis was performed for all taxa together grouped by subclade, the first including all species from the Leiocalycina + Vulpina + Red hair subclades, the second between the Microcalyx grade + Leiocalycina + Redhair subclades and the third between the Bourgonii + Microcalyx grade + Red Hair subclades (selected subclades shown in Supplementary Fig. S1; subclade selection described in Supplementary Methods, available on Dryad). We used all non-‘Fast clade’ species as outgroups to minimise the effect of substitution rate variation on *f _dM_* estimates. For each subclade subset, we calculated *f _dM_* using nonoverlapping windows of 50 informative SNPs with a rolling mean of one window. We took the absolute values of all *f _dM_* scores, as we were only interested in comparing the magnitude of introgression across loci, and calculated a mean *f _dM_* score per-locus for downstream analyses.

We then assessed whether each of our target-capture loci experienced positive selection (i.e. more non-synonymous nucleotide changes than synonymous changes) on at least one branch and at least one site using BUSTED (Murrell et al. 2015). We prepared our target capture alignments for BUSTED analysis by trimming non-homologous sequence fragments (i.e. intronic regions captured either side of exons), masking misaligned amino acid residues and producing codon-aware alignments using OMM_MACSE (Ranwez et al. 2011; Ranwez et al. 2021). Using these codon-aware alignments, we tested for the presence of selection in each locus across the same three subclade subsets of our *Inga ‘*Singlesp’ dataset for which *f _dM_* scores were calculated. We accounted for false positives by adjusting the selection test *P-* values output by BUSTED with a 5% FDR (false discovery rate) in R (Benjamini and Hochberg 1995). We assessed whether there were associations between locus selection result (‘under selection’ if the BUSTED FDR *P-*value < 0.05) and locus annotation (‘defence chemistry’, ‘differentially expressed’, ‘single-copy phylogenetically informative’ and ‘miscellaneous’) using χ^2^ tests in R. We also visualised associations between selection result and locus annotation using the R package *corrplot* (Wei et al. 2017).

Finally, we used analysis of covariance (ANCOVA) in R to assess whether variation in our per-locus *f _dM_* estimates was explained by interactions between three variables. The variables were locus annotation, selection result and the length of the locus alignment (to control for differences in the number of sites between loci). Response variables were square-root transformed to improve normalcy for all subsets except the Bourgonii + Microcalyx grade + Red hair subset, which was log-transformed. The heteroscedasticity of residuals was examined using the *plot()* function in R, and η^2^ effect sizes were calculated with the R package *effectsize* (Ben-Shachar et al. 2020). Box plots of per-locus *f* _dM_ estimates were generated using *ggplot2* (Wickham 2016) in R, with locus annotation and selection result as grouping variables. We performed both *f _dM_* and BUSTED analyses on all 1305 refined loci for each data subset, but several loci were filtered out both by BUSTED and *f _dM_*, and so we retained only those that were present in both analyses.

## Results

### Phylogenomic Analyses

We achieved a mean of 72.05% reference length recovery onto the *Mimobaits* bait sequences (78.99%, excluding outgroups, across 1305 loci). Overall, the ‘Singlesp’ dataset had 1.53×10^6^ sites, with a mean of 0.31% variable sites and 2.30% missing data across all loci. The PPD dataset had 1.89×10^6^ sites, with a mean of 0.30% variable sites and 3.02% missing data. Finally, the outgroup dataset had 1.44×10^6^ sites, with a mean of 0.66% variable sites and 12.45% missing data. The ‘Outgroup’ dataset comprised only of the 1044 ‘single-copy phylogenetically informative’ loci sequenced by previous studies (Koenen et al. 2020; Ringelberg et al. 2022). Heatmaps showing % recovery per locus are available in Supplementary Fig. S2, along with summaries of sites, variability and missing data per locus in Table S3, both available on Dryad.

Our ASTRAL analyses indicated that most bipartitions were well supported across the three datasets, with local posterior probabilities (LPP) >0.8 (Supplementary Fig. S3ai; Fig. S3b; Fig. S3c, available on Dryad) and a quartet score of 0.47, 0.69 and 1.36 for the single-accession-per-species, PPD and outgroup datasets, respectively. Within *Inga* we inferred 3 major, nested clades (‘Fast’, ‘Hairy’ and ‘Red Hair’) and 13 subclades within those (Fig. 1a). Interestingly, the concatenated IQTREE analysis of *Inga* recovered a nearly identical topology to the ASTRAL tree with high support (BS >90, Supplementary Fig. S3aii, available on Dryad), but displayed a different branching order of the Vulpina, Leiocalycina and Poeppigiana subclades. Paralog removal and trimming of hypervariable regions with PPD did not materially influence the resolution or topology of the ASTRAL *Inga* tree (Supplementary Fig. 3c, available on Dryad). Within the outgroup tree, most generic splits are well supported (LPP >0.9), and *Inga* was monophyletic. However, three *Zygia* species clustered with other genera (*Z. inundata* and *Z. sabatieri* with *Inga; Z. ocumarensis* with *Macrosamanea* (Supplementary Fig. S3b, available on Dryad)).

**Figure 1.**
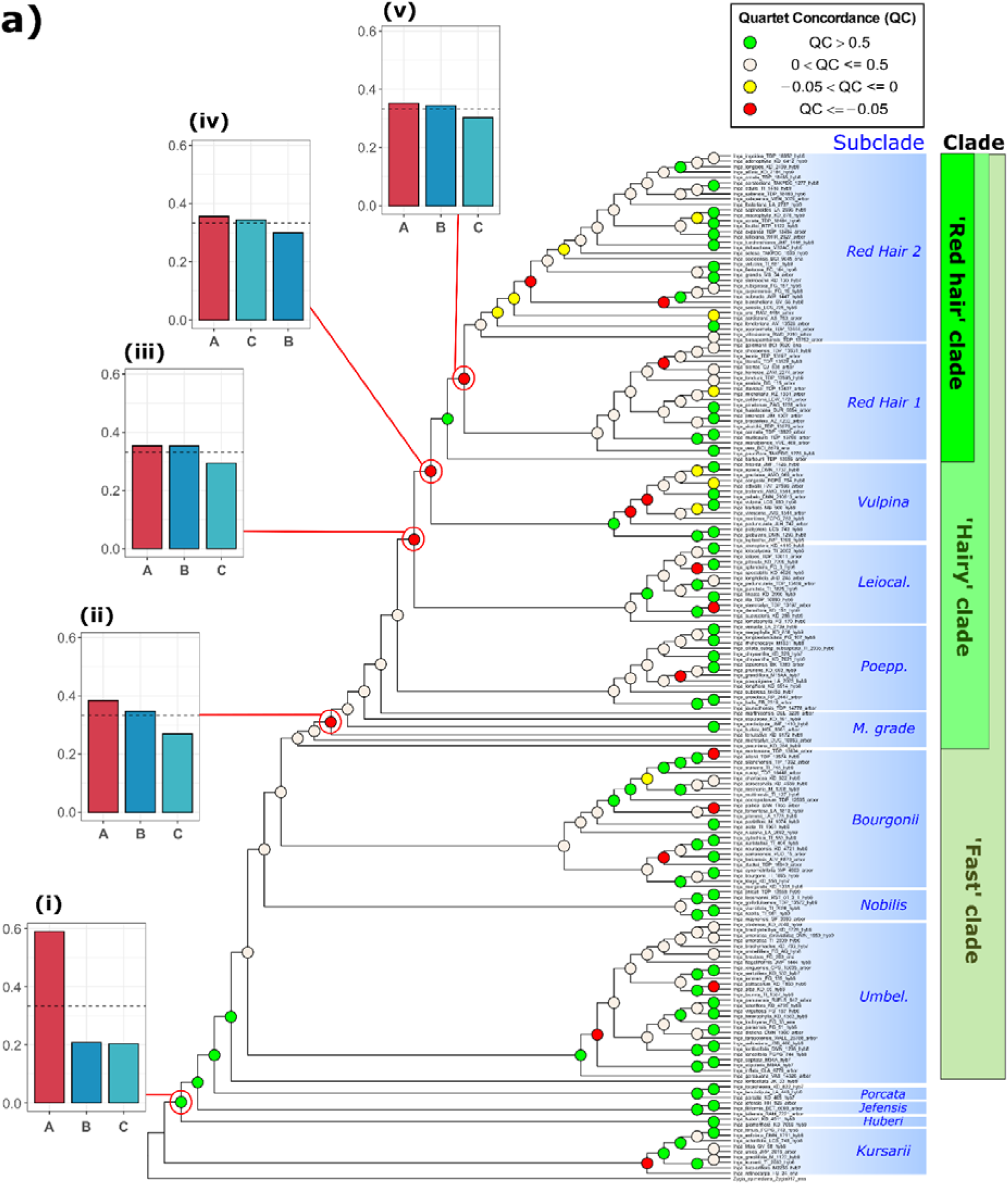
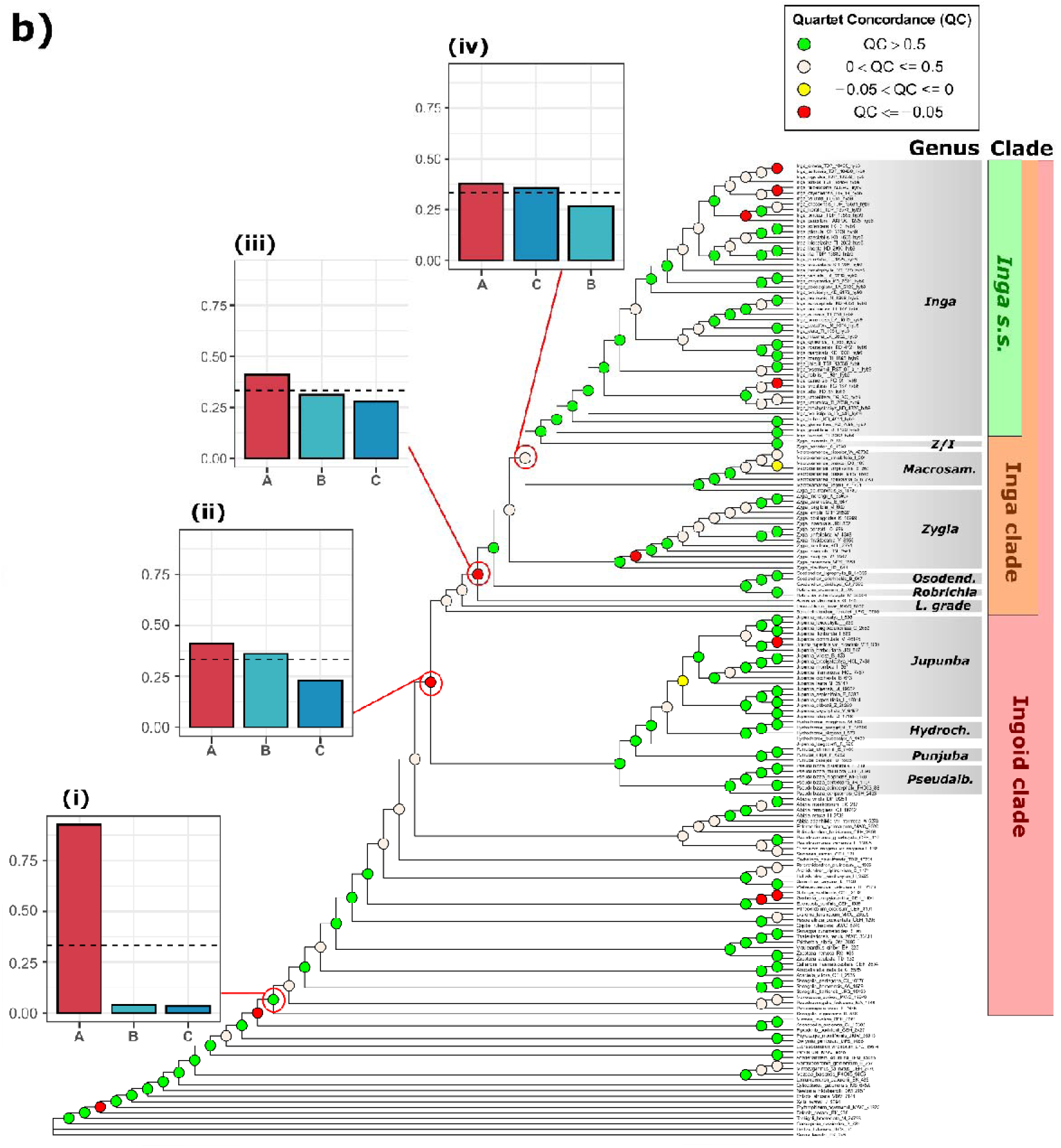
**a:** *Inga* single accession per species ASTRAL tree with QC values plotted on each node. Nodes of interest are additionally annotated with DiscoVista plots, showing the relative proportions of different discordant topologies. Quartet frequencies are represented as bar graphs, with red bars (left) representing the main topology from the ASTRAL analysis, and with blue and turquoise bars (middle and right) representing alternative topologies. Dashed horizontal lines mark the expectation for equal frequencies of the three possible topologies (Y = 0.333), i.e. maximal gene tree conflict. Node i indicates low proportions of both conflicting alternative topologies. Nodes ii-v indicate one major conflicting topology. Clades are annotated first by intrageneric subclade, and then with the broader clades within *Inga s.s.* in which they are nested (Redhair clade, Hairy clade, Fast clade). In shortened subclade annotations, ‘Leiocal.’ = Leiocalycina subclade, ‘Poepp.’ = Poeppigiana subclade, ‘M. grade’ = Microcalyx grade, ‘Umbel.’ = Umbellifera subclade. **b:** *Inga* outgroup ASTRAL tree with QC values plotted on each node. Nodes of interest are additionally annotated with DiscoVista plots, showing the relative proportions of different discordant topologies. Quartet frequencies are represented as bar graphs, with red bars (left) representing the main topology from the ASTRAL analysis, and with blue and turquoise bars (middle and right) representing alternative topologies. Dashed horizontal lines mark the expectation for equal frequencies of the three possible topologies (Y = 0.333), i.e. maximal gene tree conflict. Node i indicates low proportions of both conflicting alternative topologies. Nodes ii - iv indicate one major conflicting topology. Clades are annotated by genus, and then by the broader phylogenetic clades in which they are nested (Inga clade, Ingoid clade). In shortened genus annotations, Z/I = *Zygia/Inga*, *Macrosam.* = *Macrosamanea, Osodend. = Osodendron, L. grade = Leucochloron* grade, *Hydroch.* = *Hydrochorea. Pseudoalb. = Pseudoalbizia*.

Genetic variation visualized with SPLITSTREE showed many shared splits within *Inga, Zygia* and *Macrosamanea,* as well as between genera (Supplementary Fig. S4a, Fig. S4b available on Dryad). Shared splits are particularly evident within the ’Hairy’ clade of *Inga*, along with the ’Red hair’ clade nested within it (Supplementary Fig. S4a, available on Dryad). SPLITSTREE analysis of the Outgroup dataset showed that *Zygia inundata* and *Z. sabatieri* clustered with *Inga,* while *Z. ocumarensis* clustered with *Macrosamanea* (Supplementary Fig. S4b, available on Dryad). Paralog removal and hypervariable site trimming with PPD resulted in a different clustering of *Inga* species, with the long branch leading to *Inga gereauana* bisecting the Red hair and Hairy clades (Supplementary Fig. S4c, available on Dryad).

### Incongruence is Common Within and Among Genera

Our analyses recovered phylogenetic incongruence both within and between Ingoid clade genera (Fig.1a; Fig. 1b). Within *Inga*, negative Quartet Concordance (QC) and low Quartet Differential (QD) scores, alongside DISCOVISTA, suggested a single conflicting topology was disproportionately represented at the base of the Microcalyx grade, Leiocalycina, Vulpina, and Red hair subclades (QC in Fig. 1a nodes ii-v; QD in Supplementary Fig. S6a, available on Dryad). However, most nodes in the *Inga* ASTRAL tree showed multiple conflicting topologies in similar proportions (i.e., QC scores between 0 and 0.5; Fig. 1a). Paralog removal and trimming with PPD had minimal effect overall, but led to slightly higher QC and QD scores at some nodes (QC: Supplementary Fig. S5; QD: Fig. S6c, available on Dryad). Quartet Informativeness (QI) scores indicated all nodes were informative.

Most nodes in the outgroup tree recovered higher QC scores, indicative of more phylogenetic concordance (Fig. 1b). This was with the exception of a few nodes involving *Zygia, Macrosamanea*, *Abarema* and the clade containing *Jupunba,* that had negative QC scores (Fig. 2a nodes ii-iv; QD trees in Supplementary Fig. S6b, available on Dryad) or showed multiple conflicting topologies (QC between 0 and 0.5). For both trees, the deepest divergences had more concordant quartet topologies (Fig. 1a node i; Fig. 1b node i).

**Figure 2.**
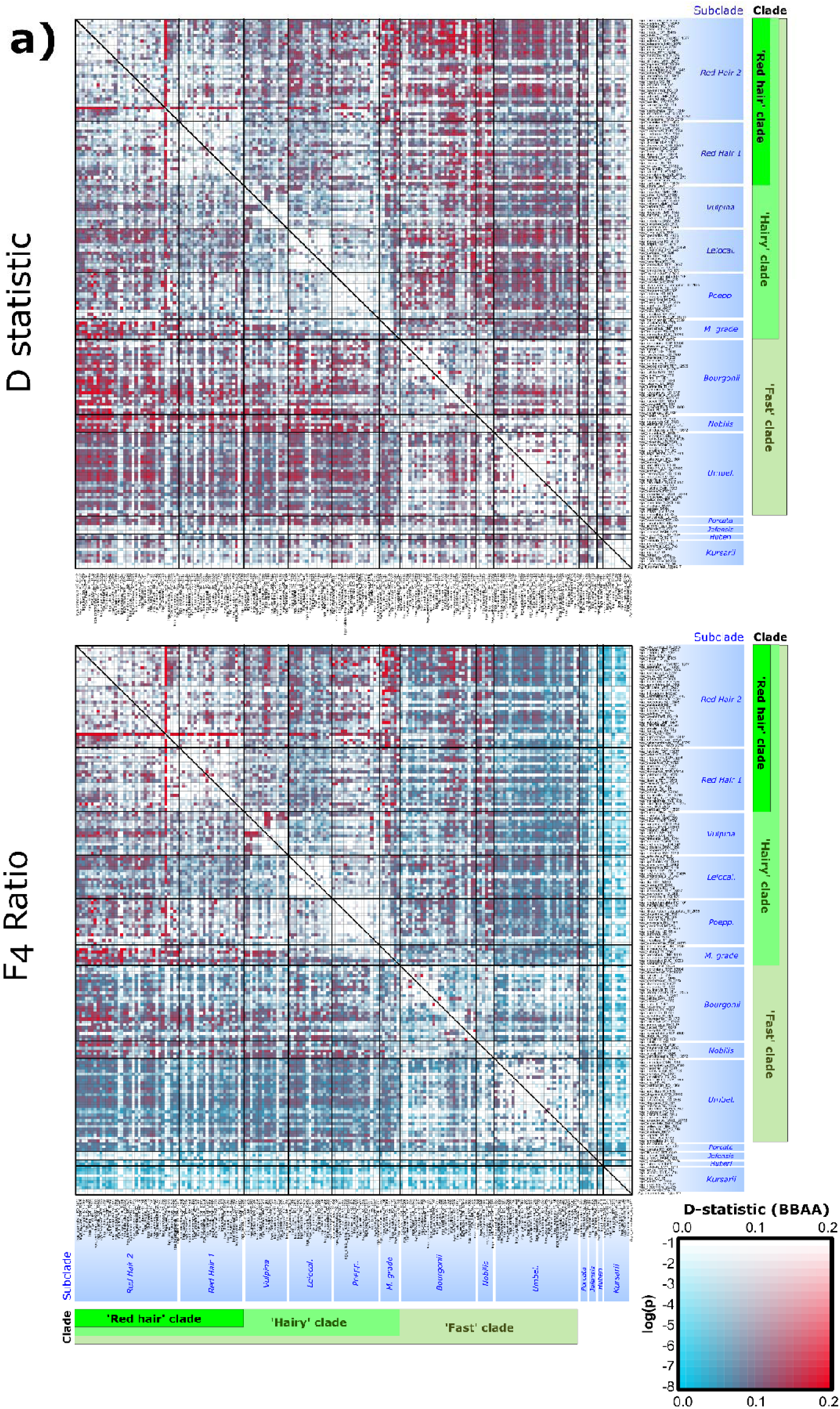
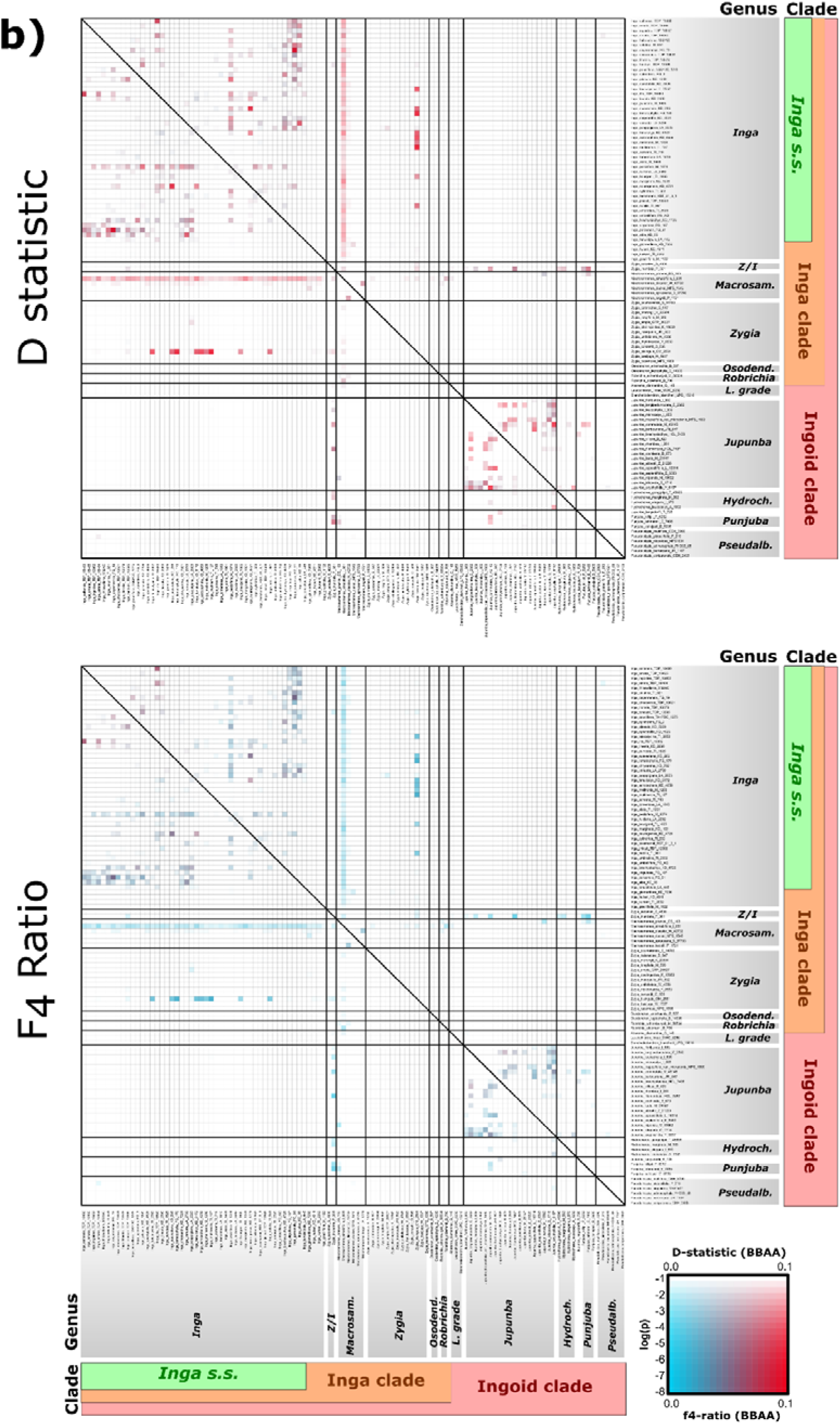
**a**: Heatmaps of per-triplet D statistic (D, above) and F_4_ ratios (below) plotted for the single-accession-per-species *Inga* dataset. Taxa P2 and P3 are displayed on the *x-* and *y*-axes in the same order as in Figure 1a. The colour of each square signifies the *D-*statistic or F_4_ ratio estimate (blue = low estimate; red = high estimate). The saturation of these colours represents the significance for that test (see log(P) in inset box, bottom right). Clades are marked in the same colours as Fig. 1a, representing subclades within *Inga* as well as the broader ‘Fast’, ‘Hairy’ and ’Red hair’ clades of *Inga*. D and F4 ratio estimates were made from species trios ordered so that the P1 and P2 taxa possess the derived allele (the BBAA pattern) more frequently than the discordant ABBA and BABA patterns. This was to ensure that the P1/P2 taxa are more closely related to each other than to the P3 taxon and outgroup, as assumed by D statistics. Clades are annotated first by intrageneric subclade, and then with the broader clades within *Inga s.s.* in which they are nested (Redhair clade, Hairy clade, Fast clade). In shortened subclade annotations, ‘Leiocal.’ = Leiocalycina subclade, ‘Poepp.’ = Poeppigiana subclade, ‘M. grade’ = Microcalyx grade, ‘Umbel.’ = Umbellifera subclade. **b**: Heatmaps of minimum per-triplet D statistic (D, above) and F_4_ ratios (below) plotted for the Outgroup dataset. Taxa P2 and P3 are displayed on the *x-* and *y*-axes in the same order as in Figure 1b. The colour of each square signifies the *D-*statistic or F_4_ ratio estimate (blue = low estimate; red = high estimate). The saturation of these colours represents the significance for that test (see log(P) in inset box, bottom right). Clades are marked in the same colours as Fig. 1b, representing different genera and the ‘Ingoid clade’, ‘Inga clade’ and the genus *Inga s.s..* D and F4 ratio estimates were made from species trios ordered so that the P1 and P2 taxa possess the derived allele (the BBAA pattern) more frequently than the discordant ABBA and BABA patterns. This was to ensure that the P1/P2 taxa are more closely related to each other than to the P3 taxon and outgroup, as assumed by D statistics. Clades are annotated by genus, and then by the broader phylogenetic clades in which they are nested (Inga clade, Ingoid clade). In shortened genus annotations, Z/I = *Zygia/Inga*, *Macrosam.* = *Macrosamanea, Osodend. = Osodendron, L. grade = Leucochloron* grade, *Hydroch.* = *Hydrochorea. Pseudoalb. = Pseudoalbizia*.

### Reticulation Occurs at Multiple Phylogenetic Scales

The overrepresentation of one incongruent topology that we inferred for several nodes (Fig. 1aii-v; Fig. 1bii-iv) was reinforced by the high *D*-statistics, F_4_ ratios and F_branch_ scores that we calculated for all three datasets. This suggests reticulation contributed to incongruence at these nodes.

Within *Inga*, significant *D*-statistics up to 0.2 were observed most frequently between the ’Red hair’ clade and the Bourgonii/Nobilis subclades, (Fig. 2a), although significant *D*-statistics were evident across the *Inga* tree even after ABBAclustering and P-value correction. ’Red hair’ clade species shared up to 20% of their sequence variation with the Microcalyx grade and the Vulpina/Leiocalycina/Poeppigiana subclades (F4 ratio = 0.2, P<0.01). F4 ratios also strongly suggested introgression events within the Red Hair clade (involving *Inga ursi*) and Vulpina subclade (involving *I. hispida/I. barbata*). Removal of putative paralogs with PPD did not materially influence the F or D statistics we inferred (Supplementary Fig. S7, available on Dryad). F_branch_ also showed ca. 20% excess allele sharing ‘Red hair’ clade species (e.g. *Inga pauciflora, I. ursi*) and the Vulpina, Poeppigiana, Bourgonii and Microcalyx grade subclades (Supplementary Fig. S8a, available on Dryad).

In the broader outgroup dataset, many *D*-statistic tests were filtered out due to insignificant clustering of ABBA patterns (Fig. 2b). However, *D*-statistics suggested some introgression in other Ingoid clade genera (*Macrosamanea* and *Jupunba*) as well as between *Zygia/Macrosamanea* and *Inga* (*D =* 0.1; Fig. 2b). F_4_ ratio and F_branch_ scores recovered more limited evidence of introgression, with the highest scores occurring within closely related species pairs in *Inga* and *Jupunba* (Supplementary Fig. S8b, available on Dryad).

Our PHYLONETWORKS analyses suggested that four reticulation events best fit the observed quartet concordance factors within *Inga* (-loglikelihood *hmax*=4, Supplementary Fig. S9ai; Table S4, available on Dryad). We inferred reticulation firstly within the ‘Red hair 2’ subclade, between the *Inga velutina* and *I. thibaudiana* lineages, with inheritance probabilities (γ) suggesting the *I. thibaudiana* lineage contributed ca. 27% of *I. velutina*’s genetic material (Fig. 3ai; γ=0.275). The other three reticulation events occurred deeper in the tree, from the *I. microcalyx* lineage into the base of the Poeppigiana/Leiocalycina/Redhair subclades (Fig. 3aii; γ=0.112), from the Red Hair 2 subclade into the Leiocalycina subclade (Fig. 3aiii; γ=0.244) and from the Umbellifera subclade/Fast clade split into the Bourgonii subclade (Fig. 3aiv; γ=0.445).

**Figure 3.**
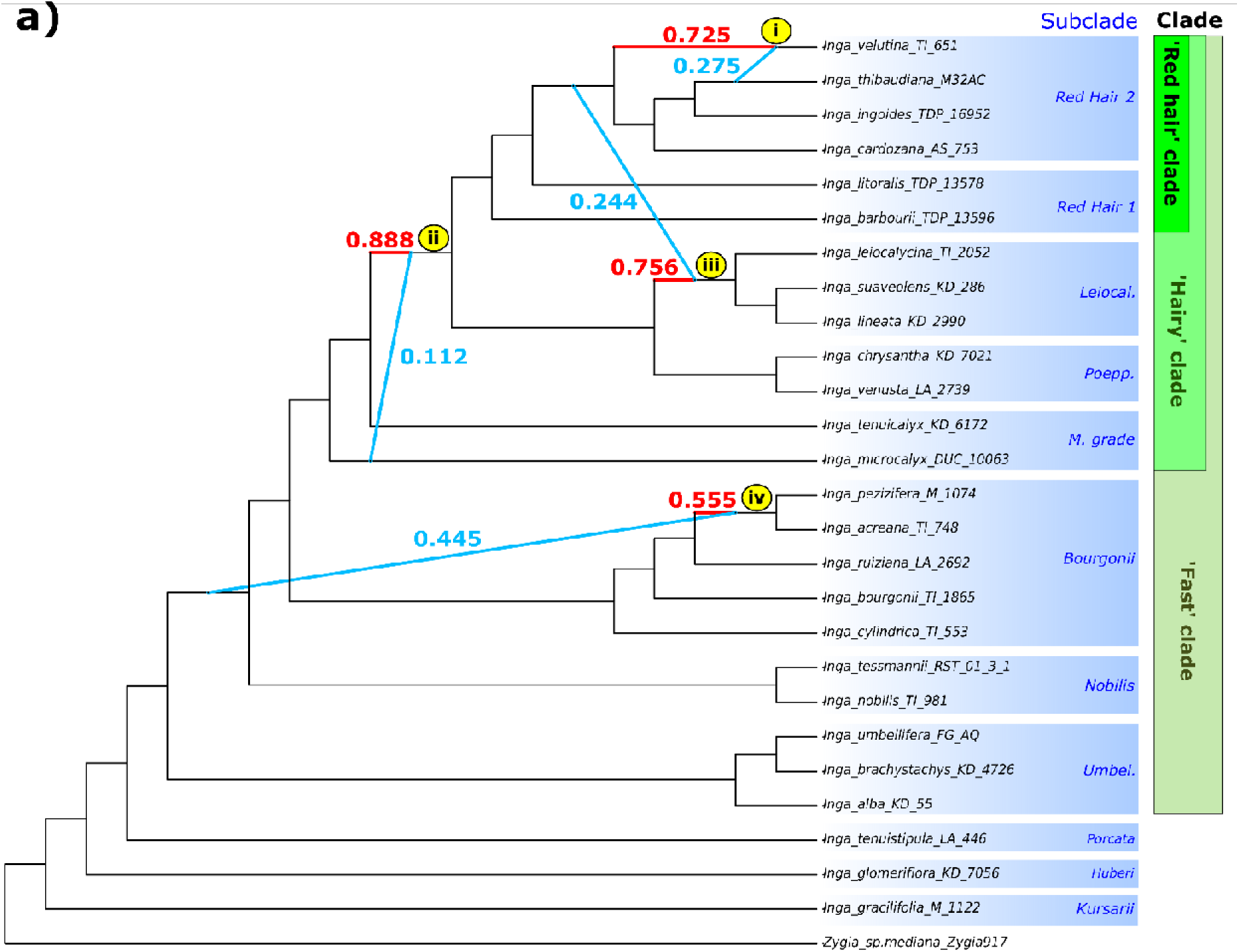
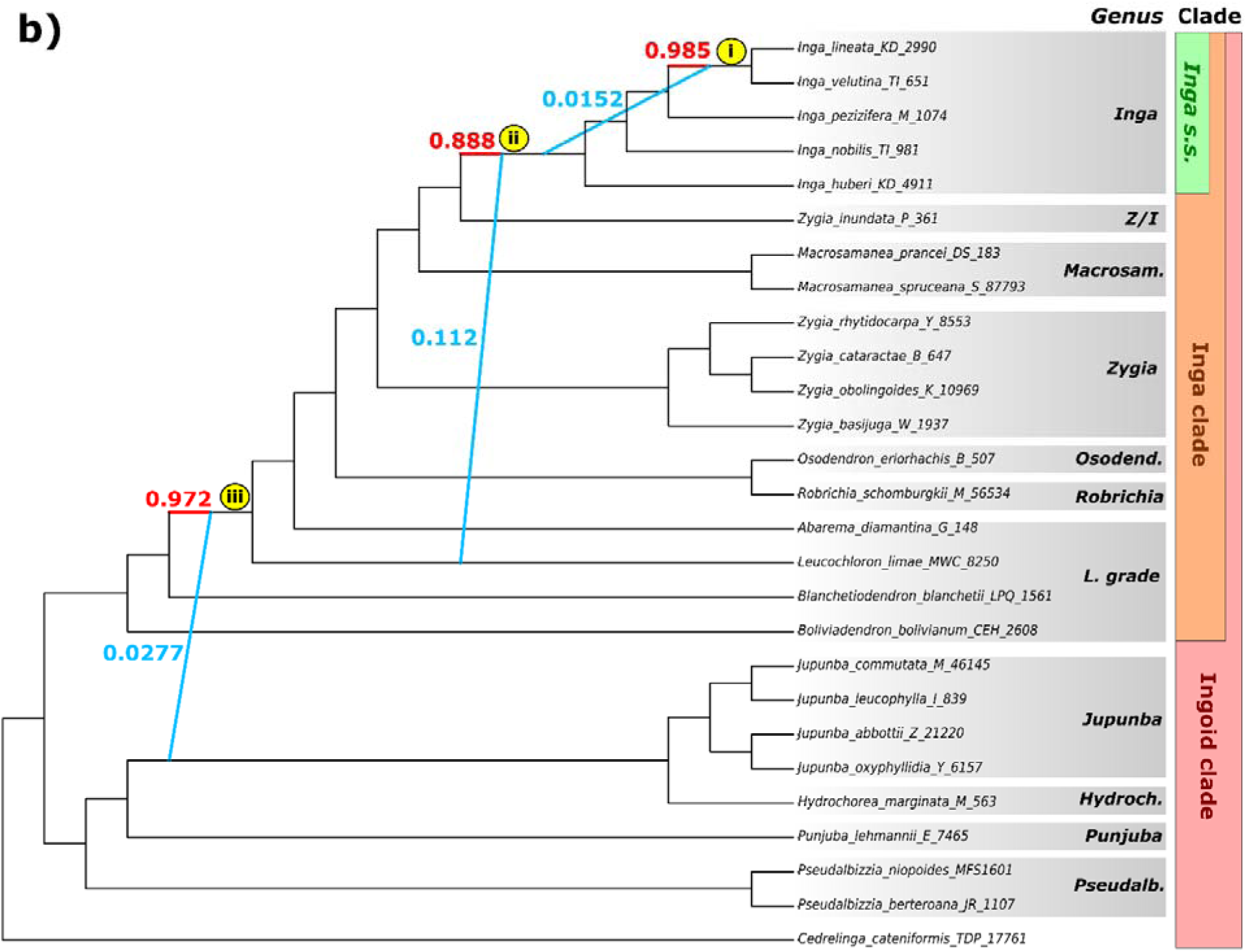
**a:** Phylogenetic network with four reticulation events (*hmax =* 4; i-iv), estimated using *SNaQ!* in the JULIA package PHYLONETWORKS. Blue and red branches indicate inferred hybridization events and numbers next to branches indicate inheritance probability (γ), roughly equal to the proportion of genetic variation contributed by each lineage to a reticulation event. Clades are annotated first by intrageneric subclade, and then with the broader clades within *Inga s.s.* in which they are nested (Redhair clade, Hairy clade, Fast clade). In shortened subclade annotations, ‘Leiocal.’ = Leiocalycina subclade, ‘Poepp.’ = Poeppigiana subclade, ‘M. grade’ = Microcalyx grade, ‘Umbel.’ = Umbellifera subclade. **b:** Phylogenetic network with three reticulation events (*hmax =* 3; i-iii), estimated using *SNaQ!* in the JULIA package PHYLONETWORKS. Blue and red branches indicate inferred hybridization events and numbers next to branches indicate inheritance probability (γ), roughly equal to the proportion of genetic variation contributed by each lineage to a reticulation event. Clades are annotated by genus, and then by the broader phylogenetic clades in which they are nested (Inga clade, Ingoid clade). In shortened genus annotations, Z/I = *Zygia/Inga*, *Macrosam.* = *Macrosamanea, Osodend. = Osodendron, L. grade = Leucochloron* grade, *Hydroch.* = *Hydrochorea. Pseudoalb. = Pseudoalbizia*.

Within *Inga* subclades, we inferred the most reticulation events within the Bourgonii subclade (*hmax*=4) and the fewest within the Vulpina subclade (*hmax=*1), with all other subclades recovering 2 reticulation events (Supplementary Fig. S9aii-aiii, available on Dryad). Many of these within-subclade events involve the same taxa with reticulate histories inferred using D statistics, F4-ratios and genus-level PHYLONETWORKS analyses (e.g. *Inga microcalyx, I. balsapambensis, I. hispida*).

Among Ingoid clade ’Outgroup’ species we inferred three reticulation events (- loglikelihood *hmax*=3, Supplementary Fig. S9b; Table S4, available on Dryad). We firstly recovered reticulation from the base of *Inga* into members of the Red Hair 2 and Leiocalycina clades (Fig. 3bi; γ=0.0152). We also inferred reticulation from the *Leucochloron limae* lineage into *Inga* (Fig. 3bii; γ=0.112) and from more distantly related Ingoid clade lineages (*Jupunba/Hydrochorea*) into the lineage leading to *L. limae* and the rest of the Inga clade (Fig. 3biii; γ=0.0277).

### Levels of Introgression and Selection Differ Between Loci in Inga

Our estimates of introgression and selection varied widely across the target capture loci we analysed. Per-locus introgression (*f _dM_*) varied between 0.0001–0.664 across all three subclade subsets, approaching the maximum *f _dM_* score of 1 in some subsets (Table 1; Supplementary Fig. S10, available on Dryad). The ‘Bourgonii + Microcalyx Grade + Red Hair’ subset produced the highest *f _dM_* scores overall (Table 1). Across the analyses, the highest *f _dM_* scores were observed in the single-copy phylogenetically informative loci, which were the most numerous (Supplementary Fig. S10, available on Dryad).

**Table 1:**
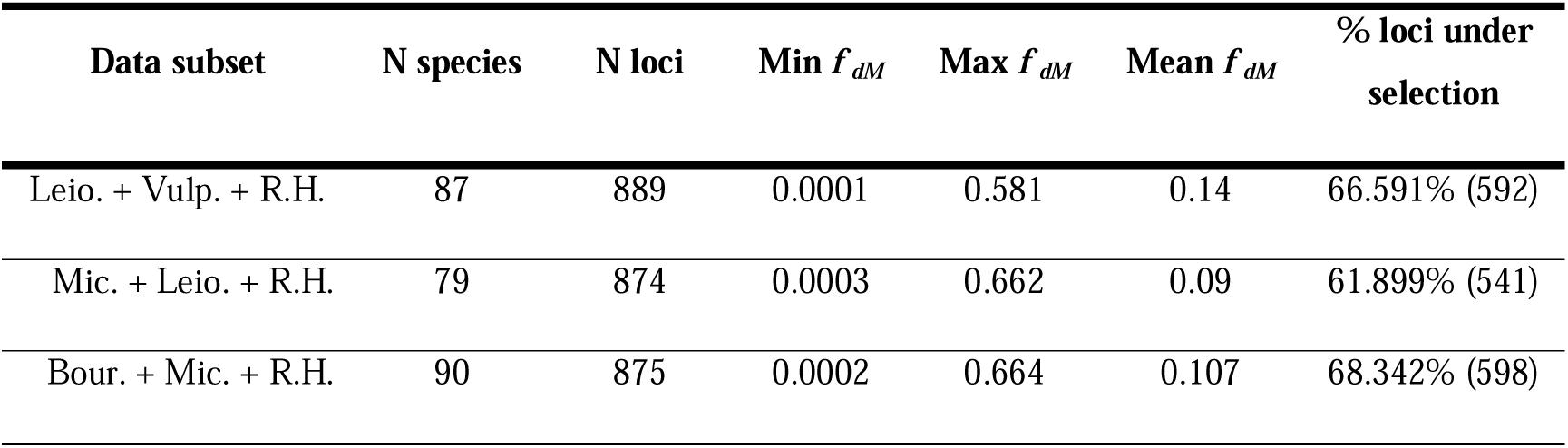
Summaries of per-locus *f* _dM_ statistics across data subsets. Columns, from left to right, indicate the subclade subset that *f _dM_* was estimated for across all loci, the number of species in each subset, the number of loci in each subset and the minimum, maximum and mean *f _dM_* scores for each subset. The phylogenetic position of the subclade data subsets that were used for running the *f _dM_*analyses listed in the leftmost column are illustrated in Supplementary Fig. S1, available on Dryad. In the first column, ‘Leio.’ = Leiocalycina subclade, ‘Vulp.’ = Vulpina subclade, ‘R.H.’ = Red Hair clade (i.e., Red Hair 1+2 subclades), ‘Mic.’ = Microcalyx grade, ‘Bour.’ = Bourgonii subclade. In the final column, the number (in parentheses) and percentage of all loci inferred to be under selection (i.e. with FDR-corrected BUSTED P-value <0.05) is shown for each subset.

In total, BUSTED inferred evidence of positive selection in between 61-68% of analysed loci per subset after multiple-testing correction (Table 1). Across all three subclade subsets, more ‘defence chemistry’ loci showed evidence of selection than the null expectation, whereas the opposite was true for other locus annotation classes (Supplementary Fig. S11, available on Dryad). However, χ^2^ tests only showed a significant association between selection result and locus annotation in the ‘Bourgonii + Microcalyx grade + Red hair’ subset (χ^2^= 10.036, df=3, N = 875, *P* = 0.0182) (Supplementary Table S5, available on Dryad).

Our ANCOVA analyses showed significant differences in *f _dM_* score means between locus annotations in the ‘Leiocalycina + Vulpina + Red hair’ subset (*P*=0.0003, *F*(3,873) *=* 6.24, η^2^ = 0.02), with ‘defence chemistry’ and ‘differentially expressed’ loci showing elevated *f _dM_* scores in loci under selection, although these were not significant (Supplementary Table S6, Fig. S10, available on Dryad). However, ANCOVA did reveal significant differences in *f _dM_* means between locus annotations when loci experienced selection in the ‘Bourgonii + Microcalyx grade + Red hair’ subset (*P*= 0.0461, *F*(3,859) *=* 5.77, ^2^ = 0.009) (Supplementary Table S6, Fig. S10, available on Dryad).

## Discussion

### Diversification of Inga and the Ingoid Clade

Our analyses recovered well-supported phylogenetic trees for *Inga* as well as the broader Ingoid clade within which it is nested (LPP >0.8, BS >90; Supplementary Fig. S3ai-ii; Fig. S3b; Fig. S3c available on Dryad). The phylogenetic tree of *Inga* we inferred marks a great advance in the resolution of inter-species relationships when compared to previous phylogenetic work using fewer loci and species (e.g. Richardson et al. 2001; Kursar et al. 2009; Dexter et al. 2010; Nicholls et al. 2015). Thus, our phylogenetic tree provides the best available framework to investigate the role of hybridisation in this species-rich group.

Our analyses revealed three nested clades within *Inga* (Fig. 1a) subdivided into twelve subclades and one grade. The deepest-level group is the ‘fast’ clade, at the base of which there was a substitution rate shift inferred by other studies (Ringelberg et al. 2023). Nested within the ‘Fast’ clade is the ‘Hairy’ clade, in which many species possess indumentum (hair-like trichomes) on young leaves as a defence against herbivores (Agrawal 1999; Coley et al. 2018). Finally, nested within the ‘Hairy’ clade is the ’Red hair’ clade, containing species that possess both dense indumentum and diverse defence chemistry (Coley et al. 2018). SPLITSTREE inferred a high degree of shared genetic variation within the ’Hairy’ and ’Red hair’ clades of *Inga*, as well as within and between other Ingoid clade genera (*Macrosamanea, Zygia* and *Jupunba*) (Supplementary Fig. S4a; S4b, available on Dryad). Of particular interest was the grouping of multiple *Zygia* species with other South American genera (e.g. *Zygia ocumarensis* with *Macrosamanea; Zygia sabatieri* and *Z. inundata* with *Inga*). The non-monophyly of *Zygia* was first described by Ferm et al. (2019), and further cases of generic non-monophyly within the Ingoid clade have been highlighted by Ringelberg et al. (2022).

### Phylogenetic Incongruence is Widespread and Influenced by Introgression

We found phylogenetic incongruence both within and between Ingoid clade genera using Quartet Concordance (QC) scores and DISCOVISTA (Fig.1a; Fig. 1b). For the Singlesp and Outgroup datasets we found one conflicting topology overrepresented at several incongruent nodes (Fig. 1a nodes ii-v; Fig 1b nodes ii-iv; see also QD trees in Supplementary Fig. S6a; Fig. S6b, available on Dryad), suggesting reticulation (Wendel and Doyle 1998). This discordance was particularly evident within the Microcalyx grade, Leiocalycina, Vulpina and Red hair subclades (Fig. 1a nodes ii-iv), subclades that also differed in branching order between the ASTRAL and concatenated IQTREE analyses (Supplementary Fig. S3ai-ii, available on Dryad). Both reticulation and incomplete lineage sorting (ILS) result in multiple evolutionary histories across the genome, and so averaging across these histories by concatenating loci often results in spurious phylogenetic relationships, explaining this difference in branching order (Degnan and Rosenberg 2009). Many deeper nodes across Figure 1a and 1b also showed several conflicting topologies at similar frequencies (QC scores between 0 and 0.5) indicating ILS, which is common in rapidly-diversifying clades like *Inga* (Degnan and Rosenberg 2009).

However, our explicit tests for introgression using *D*-statistic, F_4_ ratios and F_branch_ (Fig. 2a; Fig. 2b; Supplementary Fig. S8a, S8b, available on Dryad) also inferred widespread introgression across *Inga*. Our F_4_ ratio estimates suggested that up to 20% of genetic variation was shared between some *Inga* species, a result corroborated by the stringent F_branch_ analysis, which is less prone to inferring spurious introgression since it averages F-statistics across related branches (Supplementary Fig. S8a, available on Dryad). Interestingly, this is a similar proportion of shared variation as reported in radiations catalysed by ‘ancient’ hybridisation (e.g., Lake Victoria cichlid fish (Meier et al. 2017)). We also recovered limited evidence of reticulation in other Ingoid clade genera (e.g. *Jupunba*, *Zygia* into *Inga*) likely because most introgression signal could not be distinguished from homoplasy after stringent filtering by our ABBAclustering analysis.

PHYLONETWORKS inferred at least four migration events in *Inga* (Fig. 3a) involving several subclades with pervasive evidence of introgression in our D and F statistics (Fig. 2a). Notably, Figure 3a (node i) captures signal of the introgression events we inferred in the Red hair 2 subclade using D and F4 statistics (Fig. 2a). Similarly, nodes ii and iii of Figure 3a reflect the introgression we inferred between the Microcalyx grade and the Leiocalycina, Vulpina and Red hair subclades in our D and F statistics. Finally, the strong introgression signal we inferred in the Bourgonii and Nobilis subclades with D and F statistics (Fig. 2a) was recovered in our PHYLONETWORKS analysis (Fig. 3a, node iv), involving deep reticulation within *Inga*. We also inferred at least three migration events in the Outgroup dataset, which involved *Inga* (Fig. 3b node i-ii), the *Leucochloron* grade (node ii-iii) and *Jupunba* (node iii). While D and F4 statistics recovered most evidence of introgression in the Outgroup dataset within *Inga* and *Jupunba* (Fig. 2b), these were only broadly reflective of the PHYLONETWORKS analyses due to the stringent filtering of D and F statistic comparisons that we performed using ABBAclustering.

The widespread introgression we inferred between non-sister species throughout the radiation of *Inga* may be congruent with the syngameon hypothesis of adaptive radiation (Seehausen 2004; see also Wogan et al. 2023) rather than a ‘hybrid swarm’ preceding the radiation that catalysed diversification. Periodical hybridisation within a syngameon may be adaptive for Amazonian tree species, which are typified by large, dispersed populations (ter Steege et al. 2013). Periodical hybridisation can elevate genetic diversity and prevent Allee effects, such as inbreeding, at low population densities (Cannon and Lerdau 2015; Cannon and Lerdau 2019). *Inga* species are highly dispersible, with the entirety of Amazonia acting as a species pool for the assembly of local *Inga* communities (Dexter et al. 2017). This may facilitate introgression between *Inga* species, particularly given *Inga*’s generalist pollination syndrome and overlapping phenology (Koptur 1983). Recent work on Amazonian trees has also documented putative local syngameons in other genera, e.g. between *Brownea* species (Schley et al. 2020) and amongst three *Eschweilera* species in Brazil (Larson et al. 2021).

Further evidence for the syngameon radiation hypothesis is the introgression we inferred across whole subclades using D and F statistics (Fig. 2a; Fig. 2b; Supplementary Fig. S8a; Fig. S8b available on Dryad). Shared variation spanning subclades would be expected following introgression within ancestral syngameons, as introgressant variants are inherited by descendent species (e.g. Meyer et al. 2017; Meier et al. 2017; Schley et al. 2020). While such a pattern might result from the violation of assumptions made by D and F statistics (e.g. no substitution rate variation (Patterson et al. 2012)), we were extremely careful to account for this by using the newly implemented ‘ABBAclustering’ tool in DSUITE (Koppetsch et al. 2023). This tool tests for significant clustering of ABBA site patterns (which would be expected following introgression) to distinguish them from homoplasy (in which case ABBA patterns would be more dispersed throughout the genome).

We also inferred introgression at the base of several clades with PHYLONETWORKS (e.g. Fig. 3a, node iv; Fig. 3b, node ii), again reflecting the inheritance of introgressed loci by descendent species following ‘ancient’ hybridisation. Moreover, PHYLONETWORKS is not prone to biases caused by substitution rate variation (Koppetsch et al. 2023), and yet still recovered introgression events reflective of our *D* and *F* statistic results. Worth noting, however, is that such events may also reflect introgression with ‘ghost’ lineages that were not sampled, or have gone extinct since the introgression event (Tricou et al. 2022). This suggests introgression may be more widespread throughout *Inga*’s history than we were able to infer with our current sampling. In all, *Inga* may be representative of other large genera in neotropical rainforests, which account for half of Amazonian tree diversity. These taxa also show high sympatry in local communities, alongside emerging evidence of introgression (e.g., *Eschweilera* (Larson et al. 2021), *Protium,* (Bermingham and Dick 2001)). Assuming *Inga* is representative of these other groups, our analyses suggest that introgression is more widespread than previously thought in species-rich Amazonian tree genera (Ashton 1969).

However, incomplete lineage sorting, i.e. the retention of ancestral polymorphisms in descendent lineages (Doyle 1992), is also pervasive in rapid Amazonian tree radiations. This was shown by our results (Fig. 1a-b; Fig. 2a-2b; Supplementary Fig. 6a; Fig. S6b available on Dryad) and has been demonstrated extensively in the Mimosoid legumes, to which the Ingoid clade belongs (Koenen et al. 2020). ILS arises in these groups because the probability of coalescence (sorting of derived alleles into descendent lineages reflecting speciation history) in *t* generations decreases with increasing effective population size (Fisher 1930; Wright 1931; Kingman 2000). Most rainforest trees have large, widespread populations (ter Steege et al. 2013), such that genome-wide coalescence and sorting of alleles is unlikely to have yet occurred in rapidly-diversifying rainforest tree genera such as *Inga* (discussed in Pennington, R. T. and Lavin 2016).

### Introgression and Selection Influence the Evolution of Defence Chemistry Loci in Inga

We detected multiple deep introgression events across *Inga,* suggesting that some loci transferred by introgression are retained over time. It is likely that these loci were not immediately deleterious and were not subject to purifying selection, perhaps residing in areas of the genome that are distant from incompatibility loci, allowing them to recombine freely (Edelman et al. 2019). It is also possible that these regions are adaptive and so are maintained by positive selection, as shown in temperate tree species (e.g. Rendón-Anaya et al. 2021). This is particularly interesting in the context of chemical defences against insect herbivores, since these defences are critical for survival, co-existence and ecological divergence in *Inga* (Kursar et al. 2009; Coley et al. 2018; Forrister et al. 2023).

We found differences in the proportion of loci under selection between locus annotation classes, with elevated numbers of defence chemistry loci under selection (Supplementary Fig. S11, available on Dryad), particularly in the ‘Bourgonii + Microcalyx grade + Red hair’ subset (χ^2^ = 10.036, N= 875, df = 3, *P*-value = 0.0186) (Supplementary Table S5, available on Dryad). Previous work using phylogenetic comparative methods demonstrated divergent evolution in defence chemical profiles among sister species of *Inga* (Forrister et al. 2023), and so our results suggest a potential mechanism underlying this divergent evolution, given the molecular evidence of positive selection in defence chemistry loci we observed. This provides an important exemplar for understanding the assembly of diverse rainforest tree communities - herbivore pressure structures tree communities and so divergent defence chemistry facilitates ecological coexistence among speciose rainforest trees like *Inga* (Kursar et al. 2009; Forrister et al. 2019).

Our estimates of per-locus introgression (*f _dM_*) varied widely across the loci we analysed and across subclade subsets (Table 1; Supplementary Fig. S10, available on Dryad). Locus annotation best explained variance in introgression (*f _dM_*) across loci in the ‘Leiocalcycina + Vulpina + Red hair’ subset (*P*=0.0003, *F*(3,873) *=* 6.245, η^2^ = 0.02), with slightly higher mean introgression for defence loci under selection (Supplementary Fig. S10, available on Dryad). Similarly, for the ‘Bourgonii + Microcalyx grade + Red hair’ subset, both locus annotation and selection result best explained *f _dM_* variation (*P*=0.0461, *F*(3,859) = 5.77, η^2^ = 0.009) (Supplementary Fig. S10; Table S6, available on Dryad) but with a relatively low effect size. *f _dM_* scores were marginally higher in defence chemistry loci that were under selection in some subsets (Supplementary Fig. S10, available on Dryad), suggesting a plausible role of introgression in generating adaptive defence chemistry phenotypes in *Inga*. Moreover, novel defence chemicals in *Inga* likely arise through combination of chemical precursors, rather than *de-novo* innovation (Coley et al. 2018). This might suggest a role for admixture in generating defence chemistry, rather than solely selective mechanisms (e.g. negative frequency-dependent selection retaining rare polymorphisms (Wright 1939)). Thus, novel combinations of defences resulting from introgression may confer resistance to different herbivore communities and facilitate colonisation of, and adaptation to, new areas with different suites of herbivores. This is similar to how introgression of wing pattern genes facilitates adaptation to local mimicry rings in *Heliconius* butterflies (The Heliconius Genome Consortium 2012). While a small proportion of elevated *f _dM_* scores may result from ILS, where a local genealogical tree in a window resembles a tree expected under introgression by chance, we averaged all per-window *f _dM_* scores across loci to reduce the impact of such outliers.

In light of the Janzen-Connell hypothesis, where higher densities of conspecifics with the same defences leads to increased mortality from herbivores (Janzen 1970; Connell 1971), possession of a rare, introgressant defence chemistry phenotype is likely to be adaptive, as fewer herbivores in the new area can overcome it. Adaptive introgression facilitating colonisation of new habitats is well known in plants (SuarezCGonzalez et al. 2016), particularly in the context of defence against herbivores (Whitney et al. 2006), and may have influenced the rapid radiation of *Inga*.

## Conclusions

Our analyses indicate that rapid Amazonian tree radiations (e.g. *Inga*) display evidence of introgression, in addition to incomplete lineage sorting. The introgression we inferred may be evidence of ‘syngameons’ of co-occurring interfertile species, which are created by dispersal-assembled local tree communities in neotropical rainforests (Dexter et al. 2017). This introgression may have influenced adaptation throughout the *Inga* radiation by transferring adaptive loci between speciating lineages. Specifically, we found that loci relating to defence chemistry show more evidence of selection than expected by chance, and those loci under selection have slightly higher proportions of introgression. This suggests that introgression may facilitate adaptation, local coexistence and diversification in Amazonian trees.

## Supplementary Material

Supplementary material is available from the Dryad Digital Repository: https://doi.org/xxxxxx

## Funding

This work was supported by a Natural Environment Research Council standard grant (grant number NE/V012258/1). Sequencing was funded partly by the BBSRC, grant number BB/P022898/1. J.J.R. would like to thank the Swiss National Science Foundation Postdoc. mobility grant number P500PB_211111 for support. J.N.’s work was funded by NSF Standard and Dimensions of Biodiversity grants, numbers DEB-0640630 and DEB-1135733. This project utilised equipment funded by the Wellcome Trust (Multi-User Equipment Grant award number 218247/Z/19/Z)

## Acknowledgements

The authors acknowledge the Research/Scientific Computing teams at The James Hutton Institute and NIAB for providing computational resources and technical support for the ‘UK’s Crop Diversity Bioinformatics HPC’ (BBSRC grant BB/S019669/1), use of which has contributed to the results reported within this paper. Thanks also to Colin Hughes and Erik Koenen for their help in generating the sequencing data for the outgroup species. Many thanks to Karen Moore for her superlative help with generating the data via the Exeter Sequencing Service.

## Data availability

Data available from the Dryad Digital Repository: https://doi.org/10.5061/dryad.69p8cz92v. The accession numbers for all data collated from previous studies are found in Supplementary Table S1 on Dryad. All nucleotide sequence data produced by this study are available on NCBI GenBank under the accession numbers **XXXX**. In addition, all phylogenetic trees we produced are available on TreeBASE under the accession numbers **XXXX.**

## Notes

### Competing Interest Statement

The authors have declared no competing interest.

### Summary of Updates

Re-did D and F4 statistic analyses with ABBAclustering tool, to assess whether elevated D/F statistics are caused by introgression or homoplasy, resulting from substitution rate variation between species. Replace TreeMix analysis with PhyloNetworks, due to data violating various assumptions of TreeMix.

