## Supplementary Information for "Rampant Reticulation in a Rapid Radiation of Tropical Trees - Insights from *Inga* (Fabaceae)"

**Supplementary Figures**

- Figure S1: Clade definitions for taxon subsets used in *f _dM_* and BUSTED analyses
- Figure S2: Recovery success heatmaps
- Figure S3ai: *Inga* single-accession-per-species ASTRAL tree
- Figure S3aii: *Inga* single-accession-per-species concatenated IQtree
- Figure S3b: Outgroup ASTRAL tree
- Figure S3c: PPD ASTRAL tree
- Figure S4a: *Inga* single-accession-per-species SplitsTree
- Figure S4b: Outgroup SplitsTree
- Figure S4c: PPD SplitsTree
- Figure S5: PPD QC and DiscoVista plot
- Figure S6a: *Inga* single-accession-per-species QD tree
- Figure S6b: Outgroup QD tree
- Figure S6c: PPD QD tree
- Figure S7: *Inga* PPD D-statistic & F4 ratio plot
- Figure S8a: *Inga* single-accession-per-species Fbranch
- Figure S8b: Outgroup Fbranch
- Figure S9ai: -loglikelihood h-value plot for single-accession-per-species PhyloNetworks run
- Figure S9aii: -loglikelihood h-value plots for single-accession-per-species PhyloNetworks per-subclade run
- Figure S9aiii: Inferred networks for single-accession-per-species per-subclade run
- Figure S9b: -loglikelihood h-value plot for Outgroup PhyloNetworks run
- Figure S10: Boxplots of *f _dM_* grouped by locus class and selection result for subclade subsets
- Figure S11: Correlation plots of selection results grouped by locus class for subclade subsets

**Supplementary Tables (Available on Dryad)**

- Table S1: Voucher information
- Table S2: DiscoVista clade definitions
- Table S3: Locus recovery success and alignment summaries
- Table S4: -loglikelihood values per value of *h* for PhyloNetworks analyses
- Table S5: BUSTED selection χ2 results
- Table S6: ANCOVA BUSTED *f _dM_* results

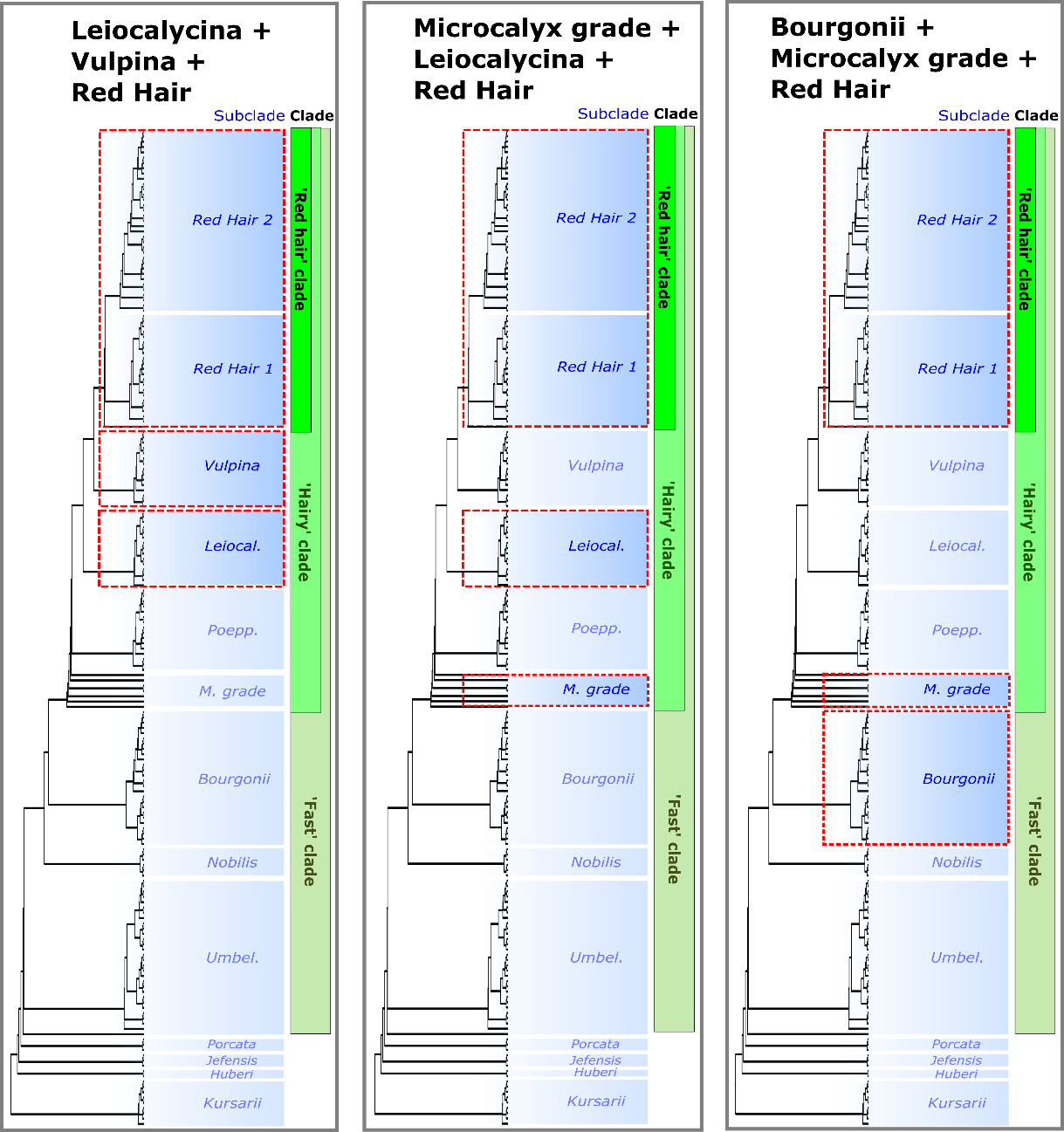

**Figure S1**: Definitions of subclades chosen for the *f _dM_* and BUSTED analyses, indicating the three data subsets chosen within *Inga* based on inference of introgression events in these groups using PhyloNetworks and Dsuite. Subclades and broader, generic-level clades are marked in the same colours as Fig. 1a.

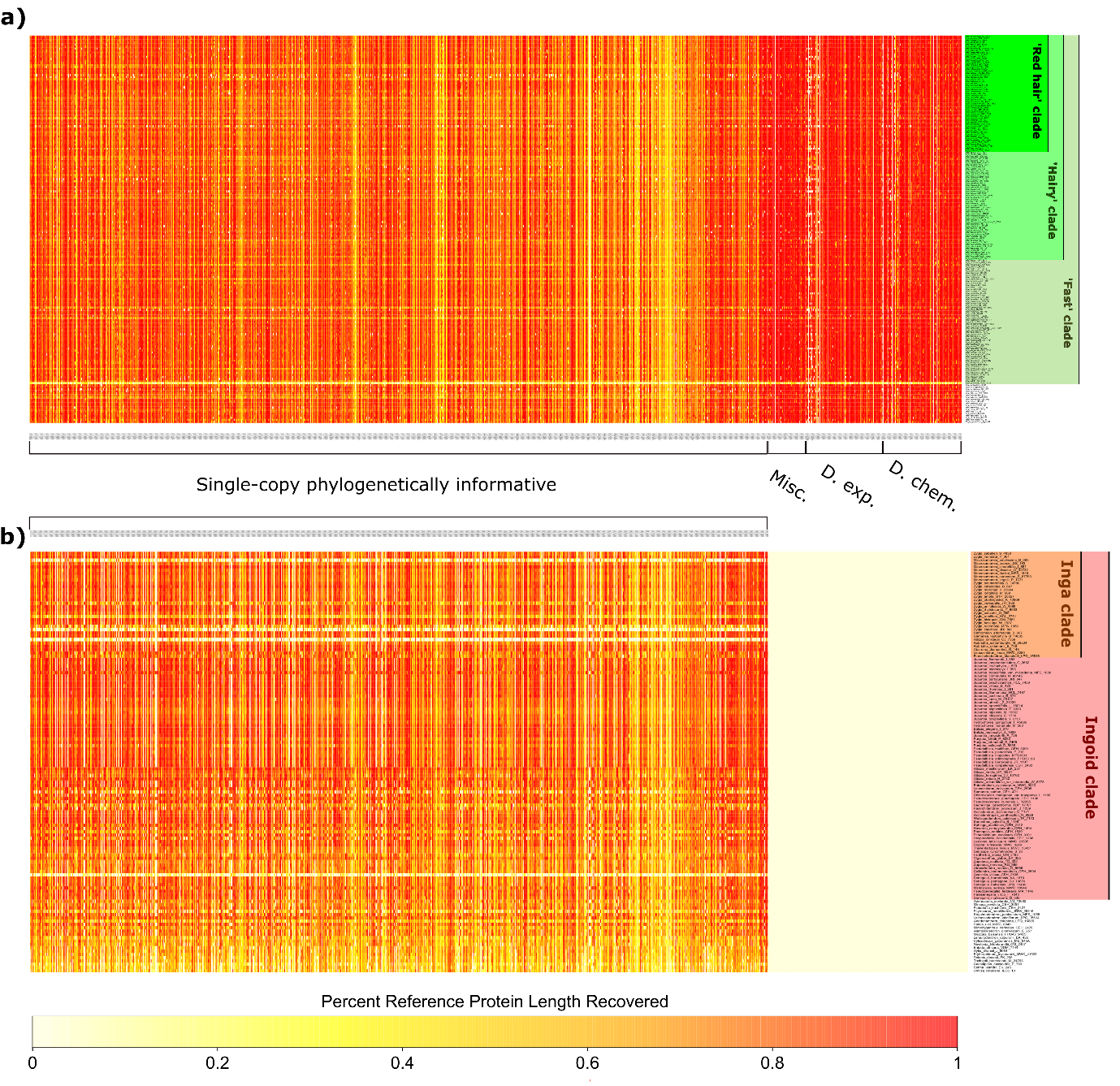

**Figure S2:** Locus recovery heatmap, indicating percentage of the reference protein length recovered for each locus in the ‘Mimobaits’ bait kit (x axis) for each accession in the phylogenomic datasets (y axis). On the spectrum, yellow indicates <20% length recovered, through to red, which reflects >80% length recovered.

a) Recovery success for the *Inga* ‘Singlesp’ dataset, including all loci from all annotation groups annotated on the x axis (Single-copy phylogenetically informative; ‘Misc.’ = Miscellaneous; ‘D. exp’ = Differentially expressed; ‘D. chem’ = Defence chemistry). Clades are marked in the same colours as Fig. 1a, representing the ‘Fast’, ‘Hairy’ and 'Red hair' clades.
b) Recovery success for the ‘Outgroup’ dataset, for which only ‘Single-copy phylogenetically informative’ loci were sequenced. Clades are marked in the same colours as Fig. 1b, representing the ‘Ingoid clade’ and ‘Inga clade’.

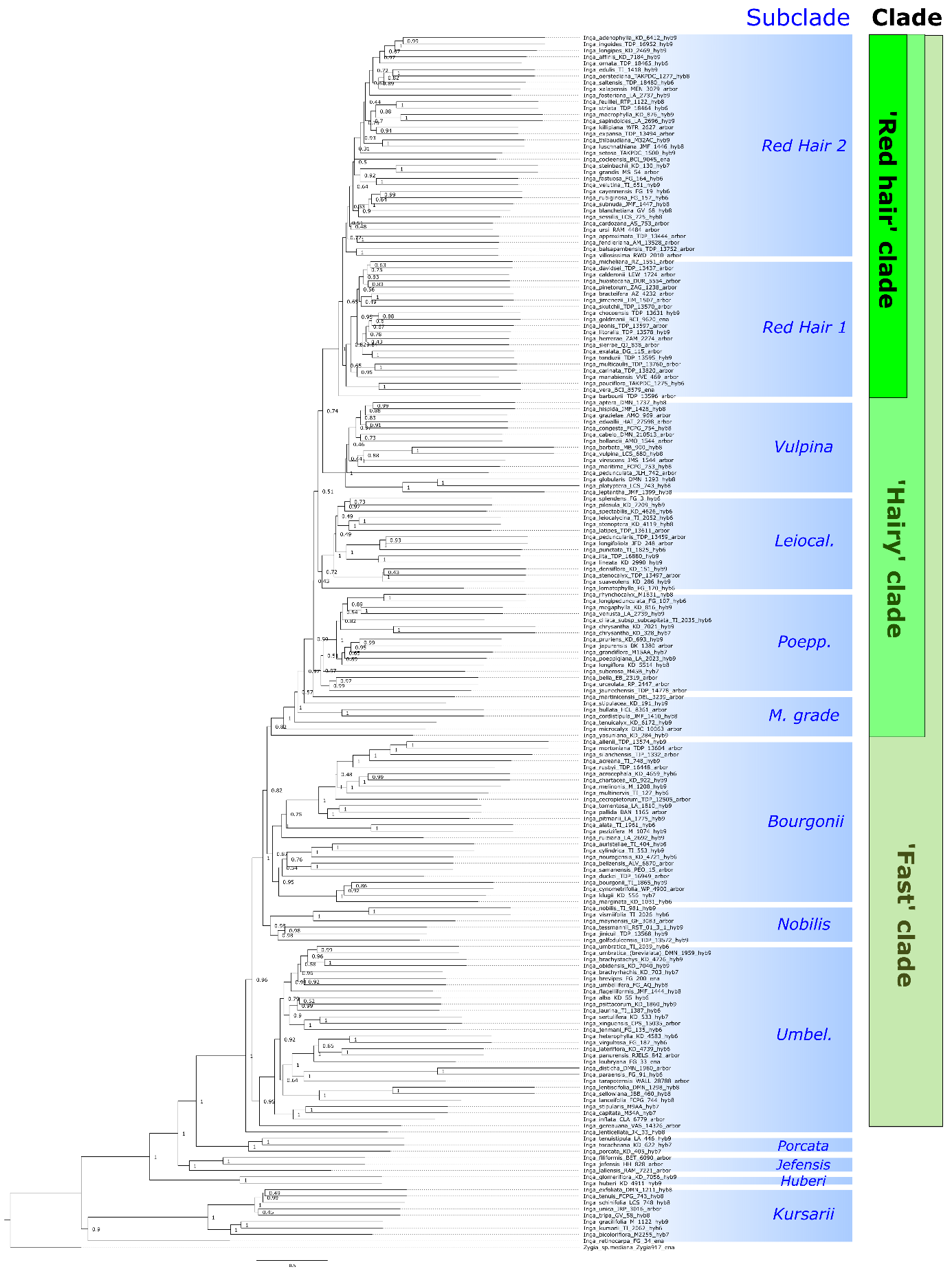

**Figure S3ai:** *Inga* single-accession-per-species (‘Singlesp’) phylogenetic tree inferred using ASTRAL. Clades and subclades are marked in the same colours as Fig. 1a. Local posterior probabilities (LPP), a measure of topological support, are shown for each node. An LPP score of 1 = full support.

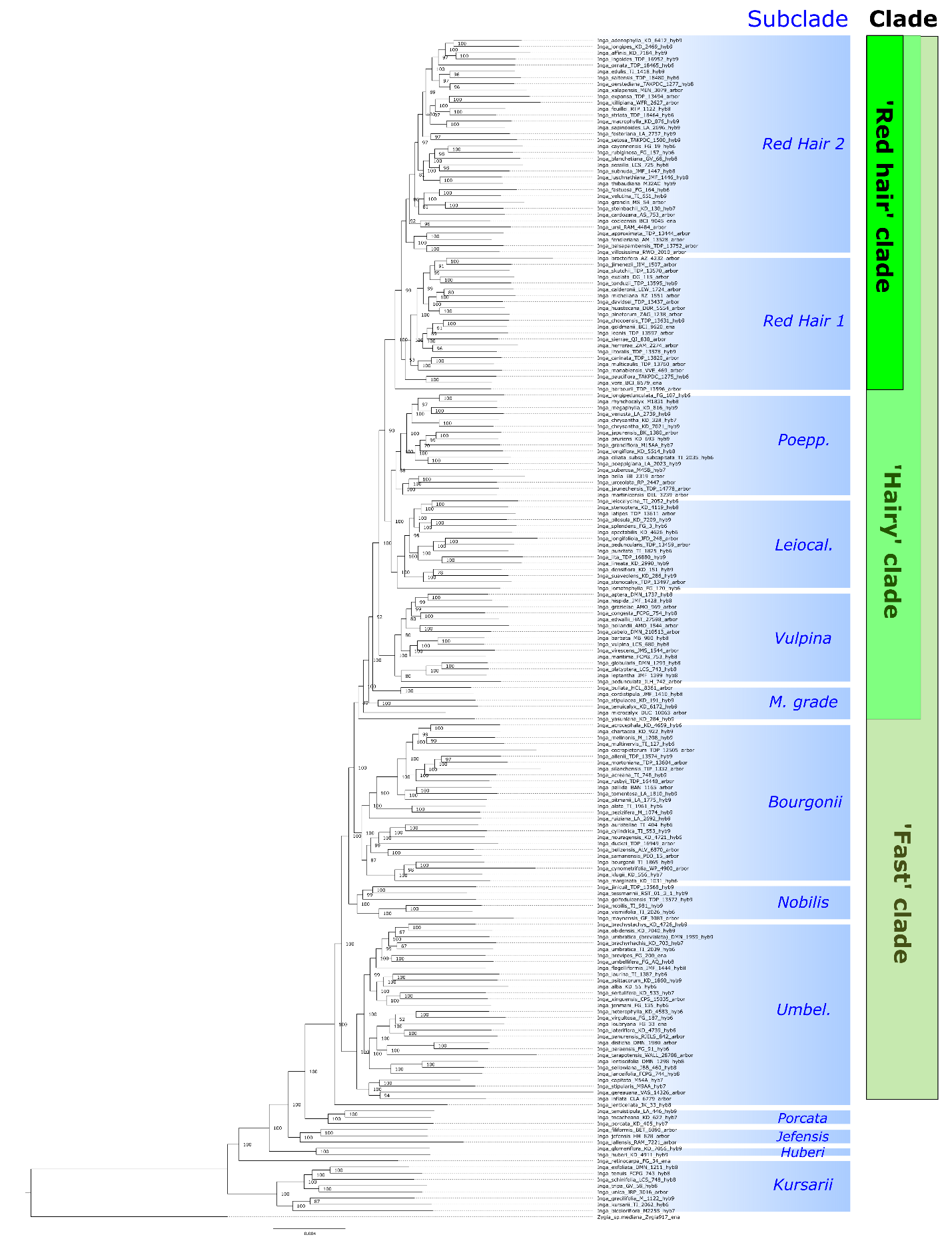

**Figure S3aii:** *Inga* single-accession-per-species (‘Singlesp’) phylogenetic tree inferred from a concatenated supermatrix of all loci using IQtree. Clades and subclades are marked in the same colours as Fig. 1a. UltraFast bootstrap support values, a measure of topological support, are shown for each node. Values >95% indicate good support for a clade.

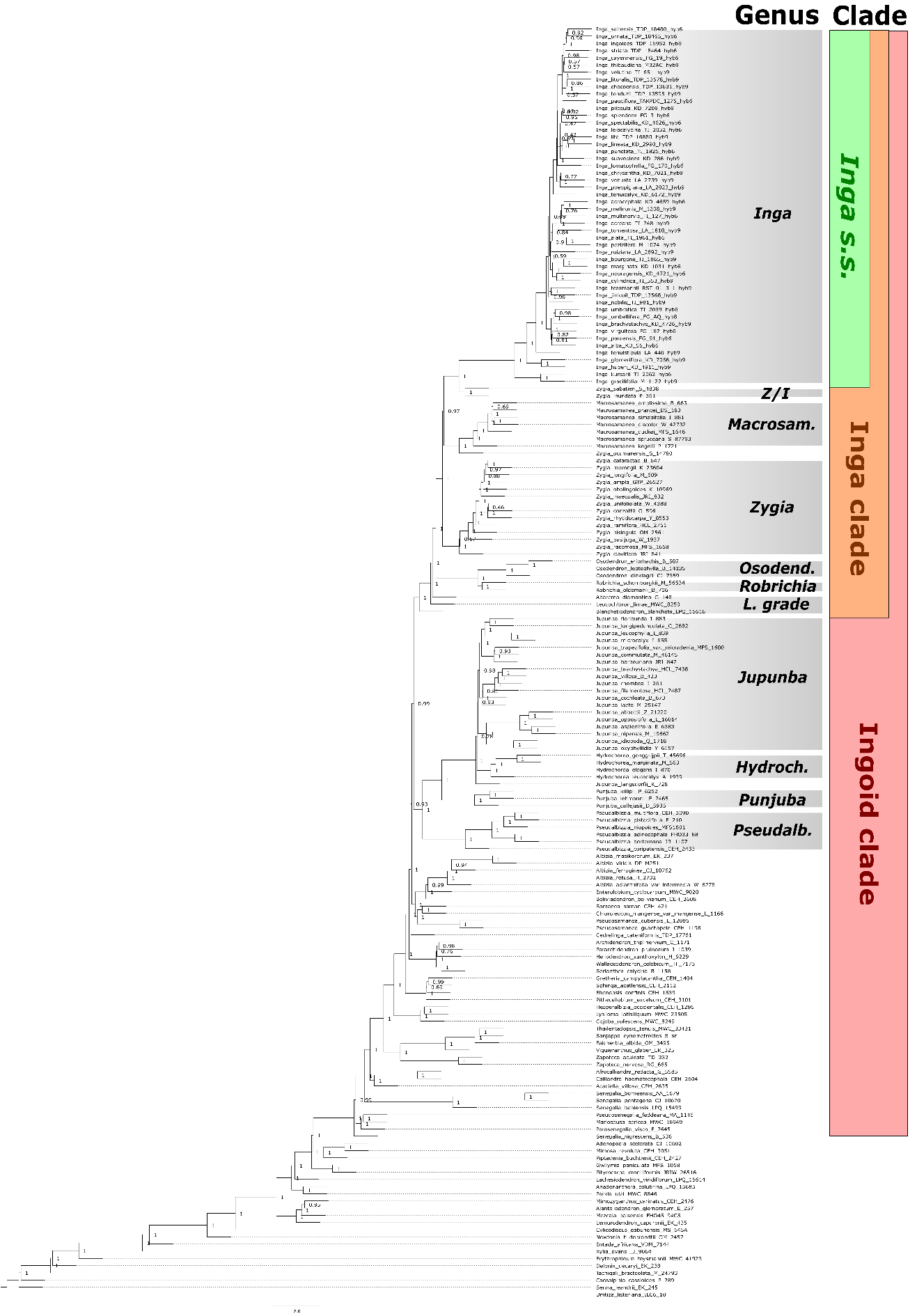

**Figure S3b:** ASTRAL phylogenetic tree inferred for the ‘Outgroup’ dataset. Clades and genera are marked in the same colours as Fig. 1b. Local posterior probabilities (LPP), a measure of topological support, are shown for each node. A score of 1 = full support.

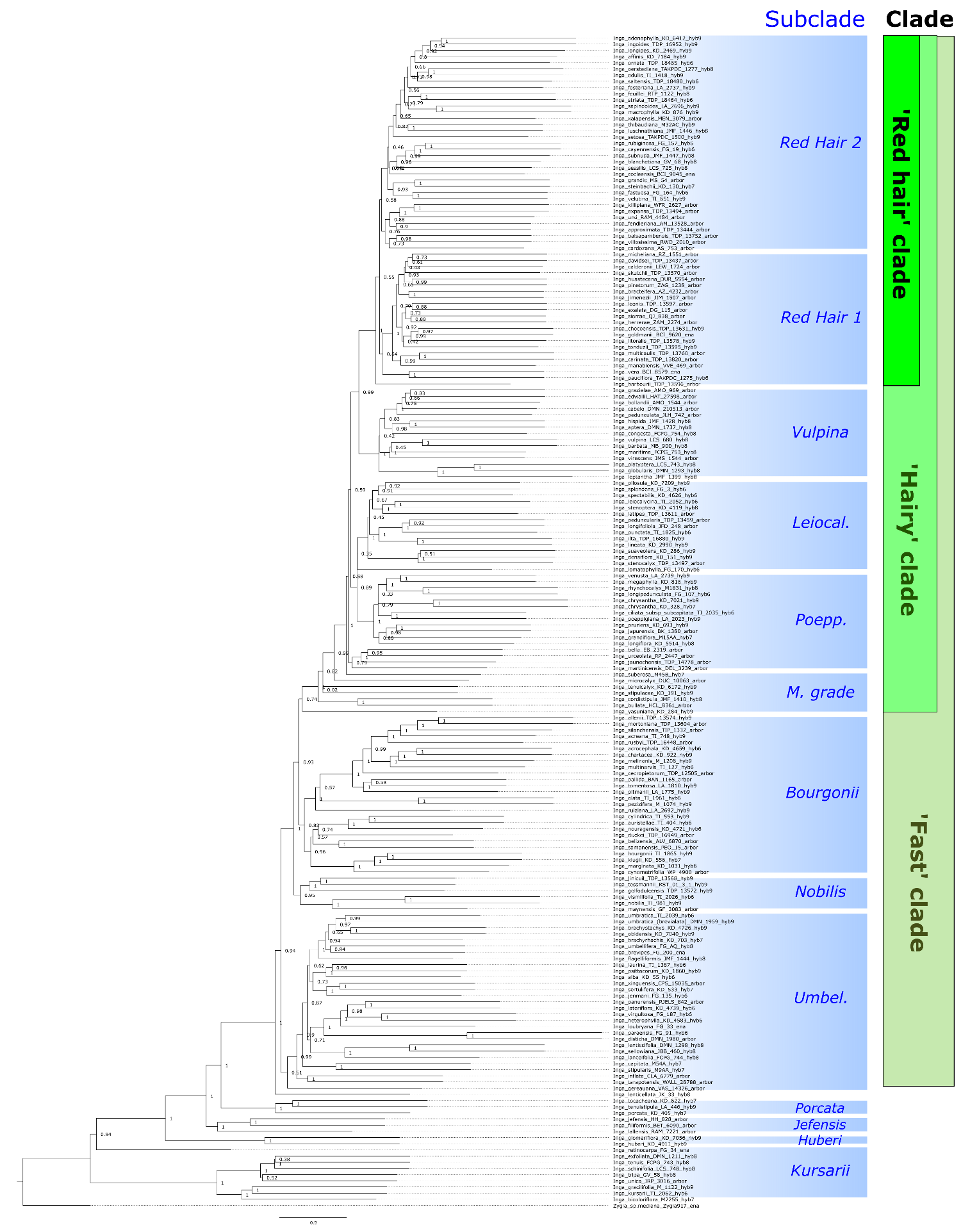

**Figure S3c:** ASTRAL phylogenetic tree inferred for the ‘PPD’ dataset with putative paralogs removed. Clades and subclades are marked in the same colours as Fig. 1a. Local posterior probabilities (LPP), a measure of topological support, are shown for each node. An LPP score of 1 = full support.

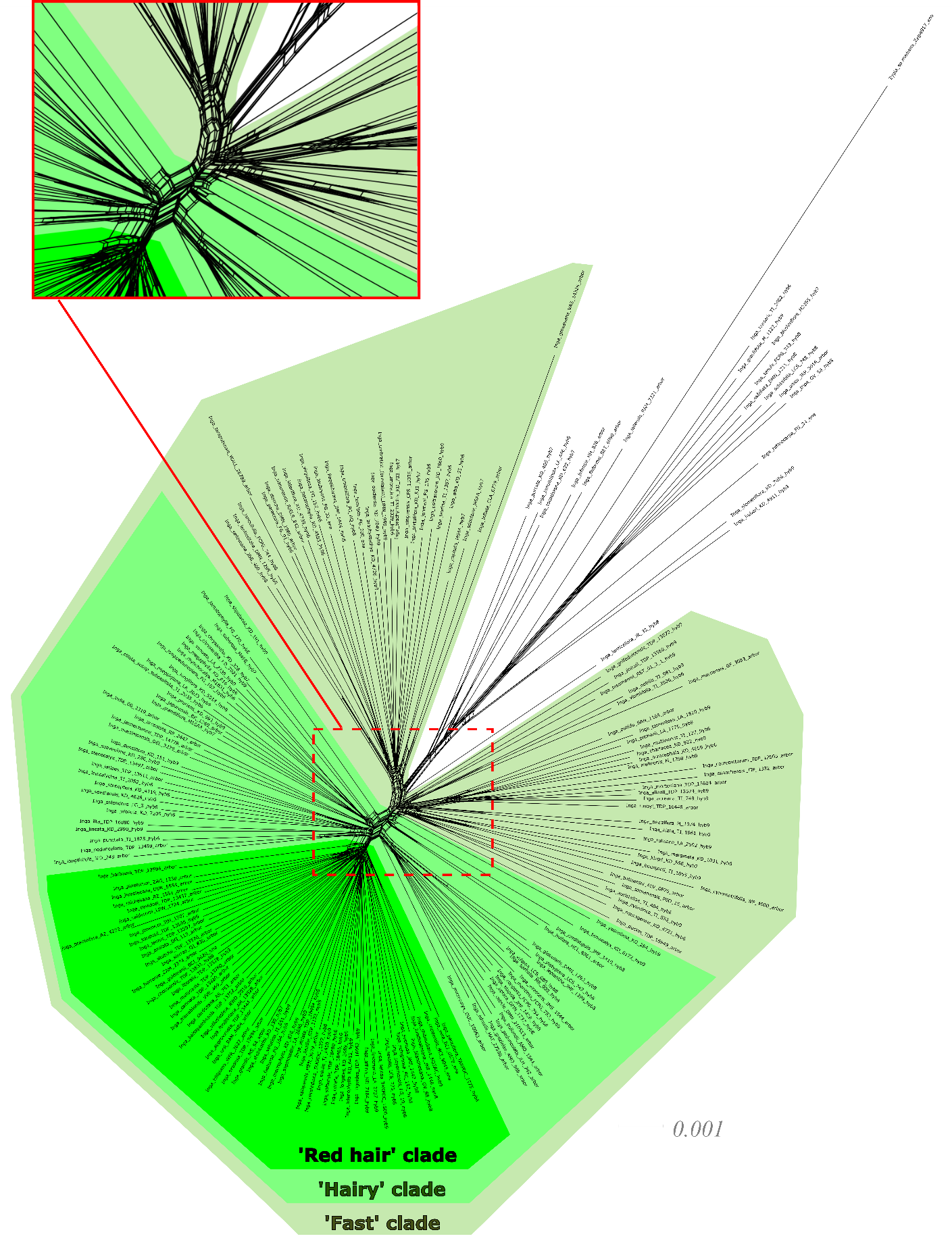

**Figure S4a:** *Inga* single-accession-per-species (‘Singlesp’) SplitsTree built using uncorrelated P distances. Zoomed-in area at the base of multiple *Inga* clades highlights many shared splits. Clades are marked in the same colours as Fig. 1a, representing the ‘Fast’, ‘Hairy’ and 'Red hair' clades.

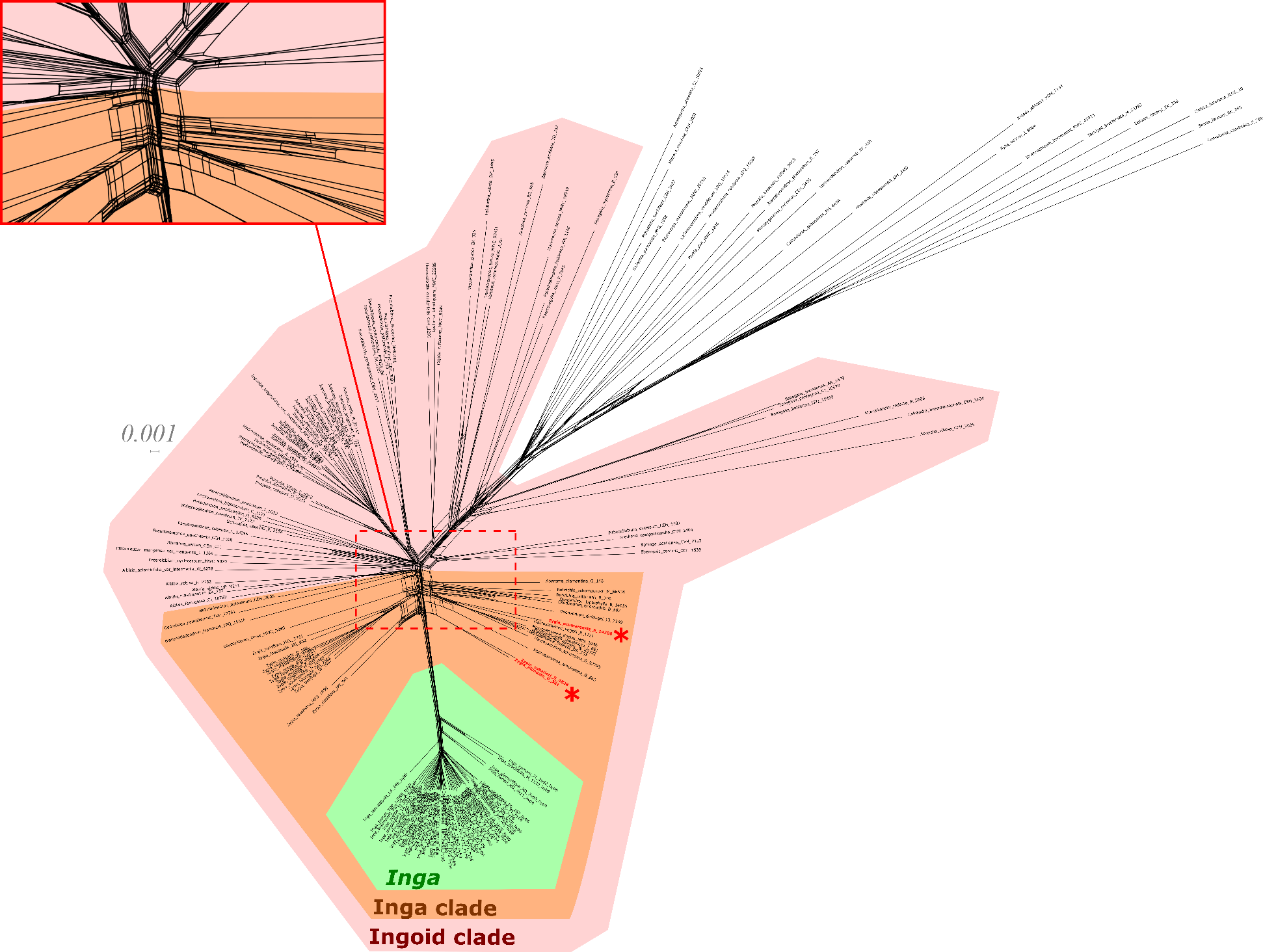

**Figure S4b:** Outgroup SplitsTree built using uncorrelated P distances. Zoomed-in area at the base of multiple genera indicates many shared splits. Species that are emboldened, marked in red and with a red asterisk are those that do not cluster with their congenerics. Clades are marked in the same colours as Fig. 1b, representing the ‘Ingoid clade’, ‘Inga clade’ and the genus *Inga*.

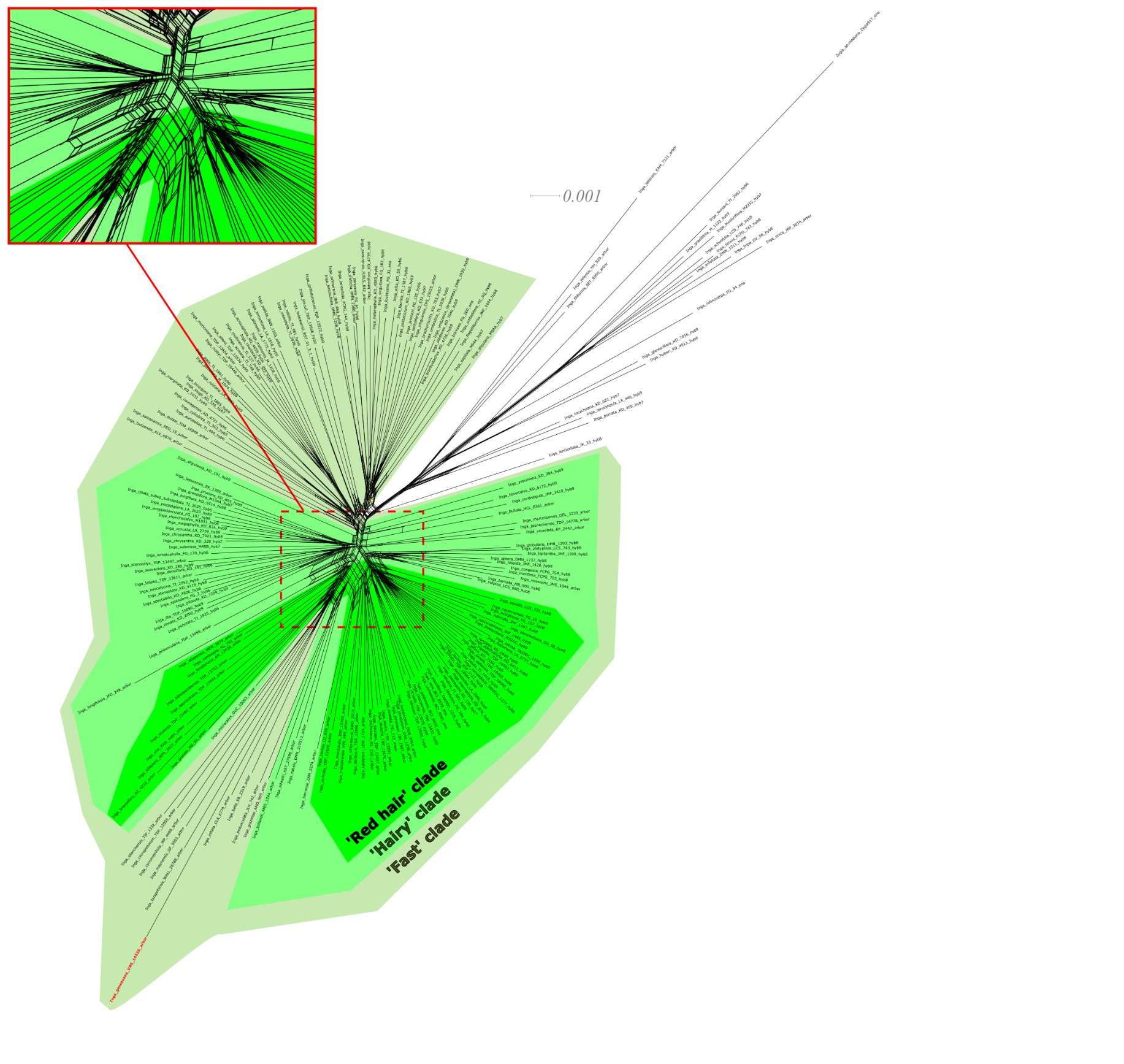
**Figure S4c:** *Inga* PPD SplitsTree built using uncorrelated P distances and putative paralogs removed. Zoomed-in area at the base of multiple *Inga* clades highlights many shared splits. *Inga gereauana* is emboldened and coloured in red due to its long branch, and the fact that it bisects both the ‘Red hair’ and ‘Hairy’ clades. Clades are marked in the same colours as Fig. 1a, representing the ‘Fast’, ‘Hairy’ and 'Red hair' clades.

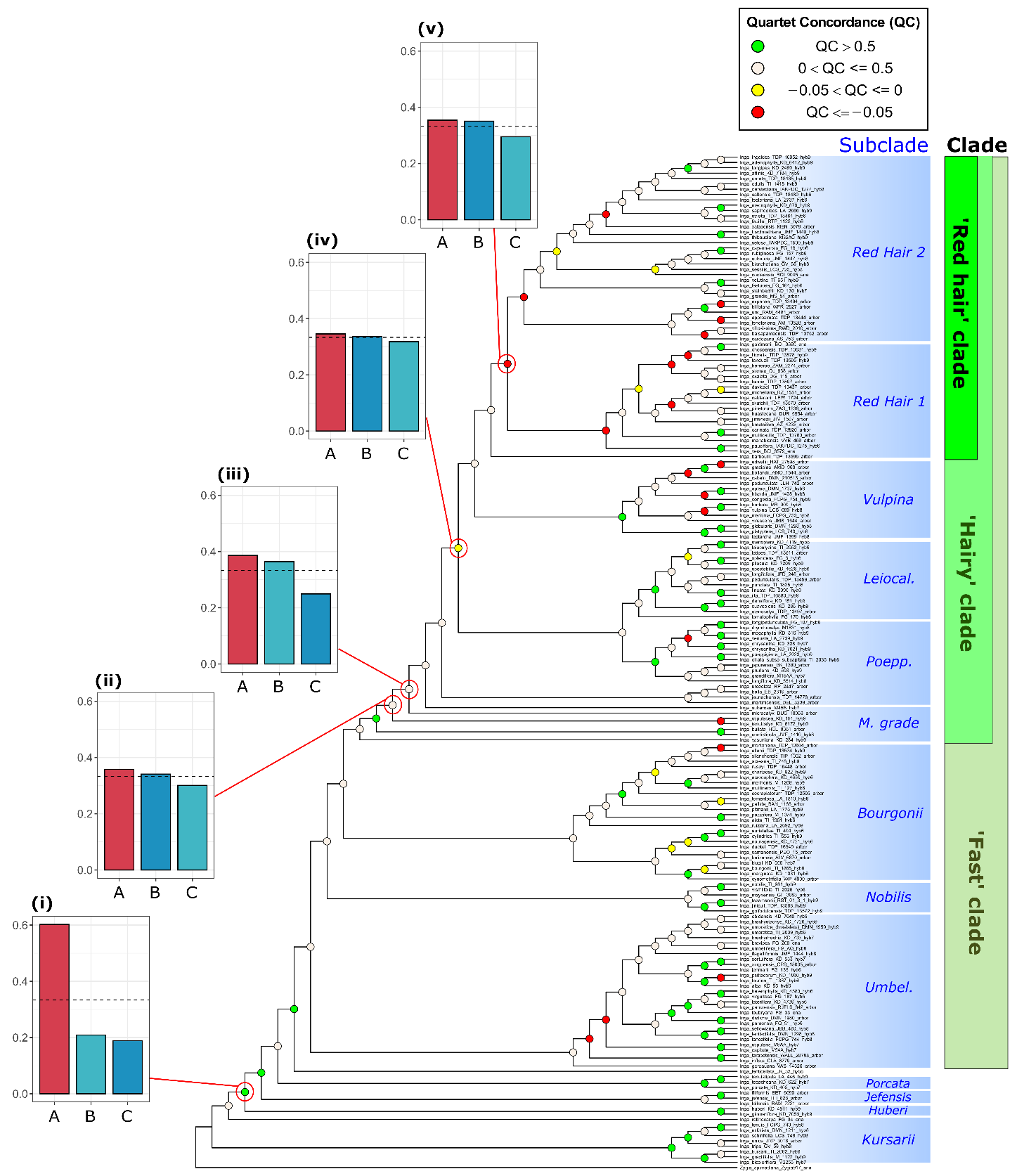

**Figure S5:** *Inga* PPD ASTRAL tree with QC values plotted on each node. Nodes of interest are additionally annotated with DiscoVista plots, showing the relative proportions of different discordant topologies. Quartet frequencies are represented as bar graphs, with red bars (left) representing the main topology from the ASTRAL analysis, and with blue and turquoise bars (middle and right) representing alternative topologies. Dashed horizontal lines mark the expectation for equal frequencies of the three possible topologies (Y = 0.333), i.e. maximal gene tree conflict. Node i indicates low proportions of both conflicting alternative topologies. Nodes ii-v indicate one major conflicting topology. Subclades and generic-level clades are labelled as in Fig. 1a.

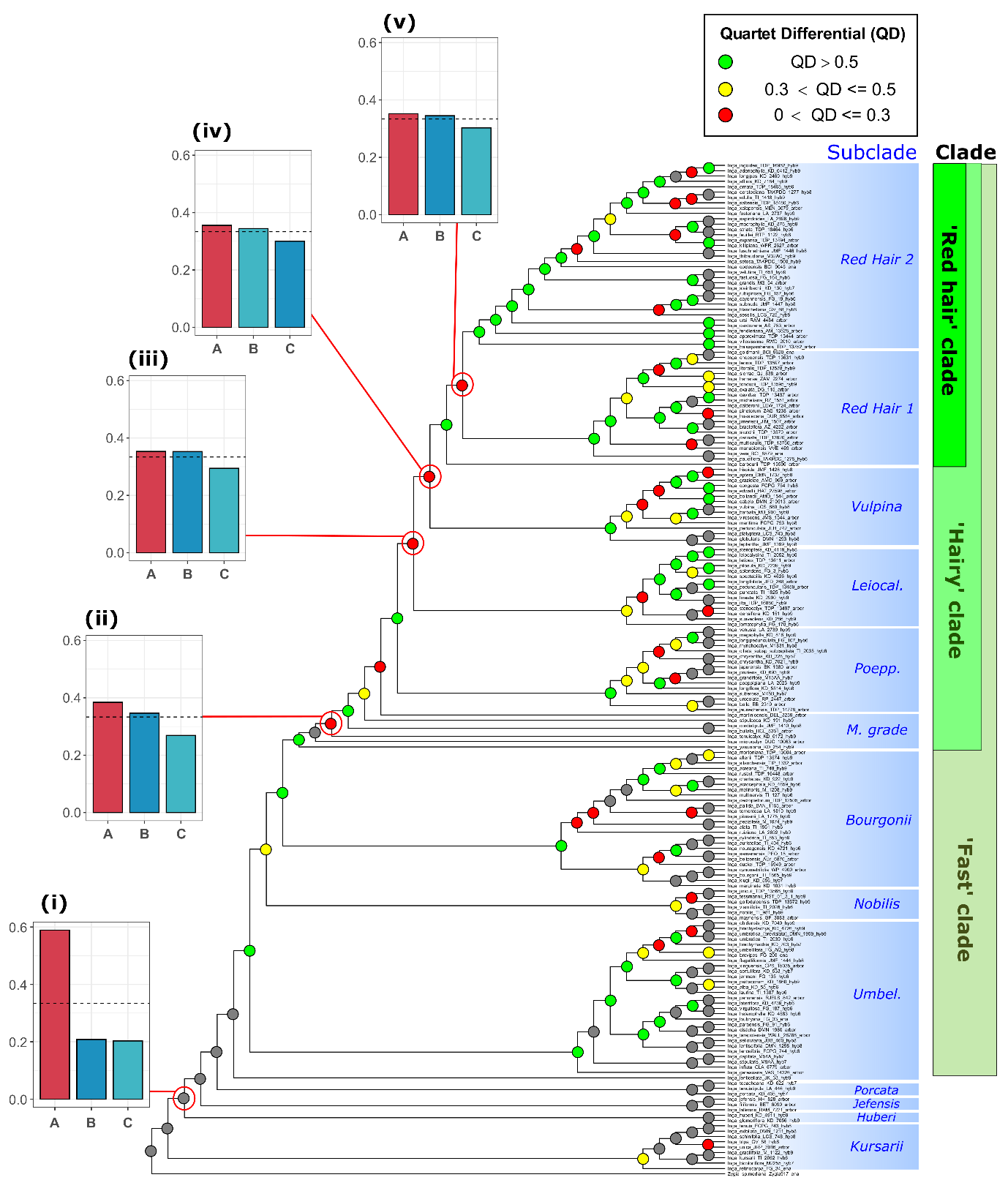

**Figure S6a:** *Inga* single accession per species (‘Singlesp’) ASTRAL tree with QD values plotted on each node, indicating the differential between the frequency of the main topology with conflicting topologies (low QD = one incongruent topology favoured). Nodes of interest are additionally annotated with DiscoVista plots, showing the relative proportions of different discordant topologies. Quartet frequencies are represented as bar graphs, with red bars (left) representing the main topology from the ASTRAL analysis, and with blue and turquoise bars (middle and right) representing alternative topologies. Dashed horizontal lines mark the expectation for equal frequencies of the three possible topologies (Y = 0.333), i.e. maximal gene tree conflict. Node i indicates low proportions of both conflicting alternative topologies. Nodes ii-v indicate one major conflicting topology. Subclades and generic-level clades are labelled as in Fig. 1a.

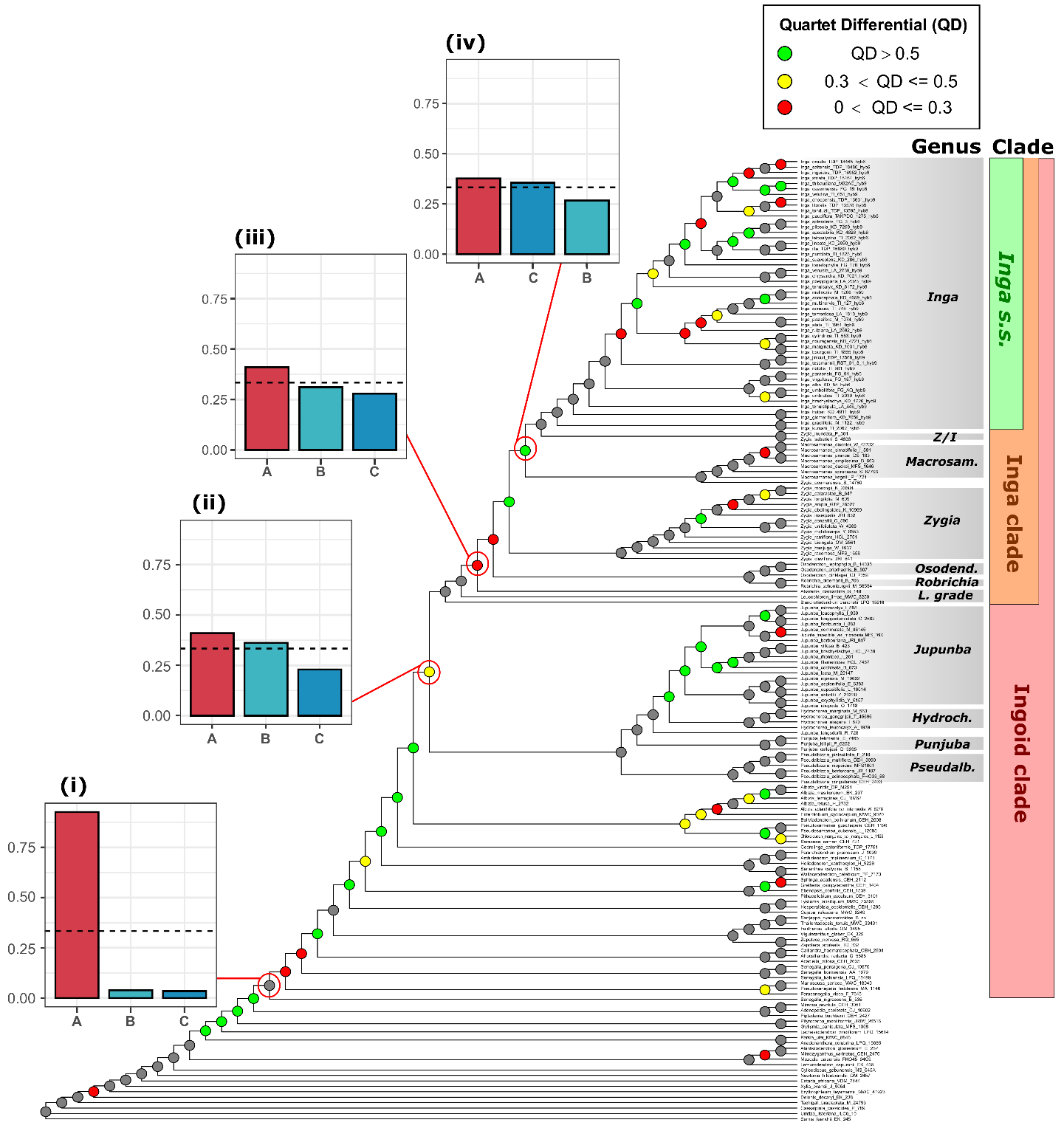

**Figure S6b:** ‘Outgroup’ dataset ASTRAL tree with QD values plotted on each node, indicating the differential between the frequency of the main topology with conflicting topologies (low QD = one incongruent topology favoured). Nodes of interest are additionally annotated with DiscoVista plots, showing the relative proportions of different discordant topologies. Quartet frequencies are represented as bar graphs, with red bars (left) representing the main topology from the ASTRAL analysis, and with blue and turquoise bars (middle and right) representing alternative topologies. Dashed horizontal lines mark the expectation for equal frequencies of the three possible topologies (Y = 0.333), i.e. maximal gene tree conflict. Node i indicates low proportions of both conflicting alternative topologies. Nodes ii-iv indicate one major conflicting topology. Genera and clades are labelled as in Fig. 1b.

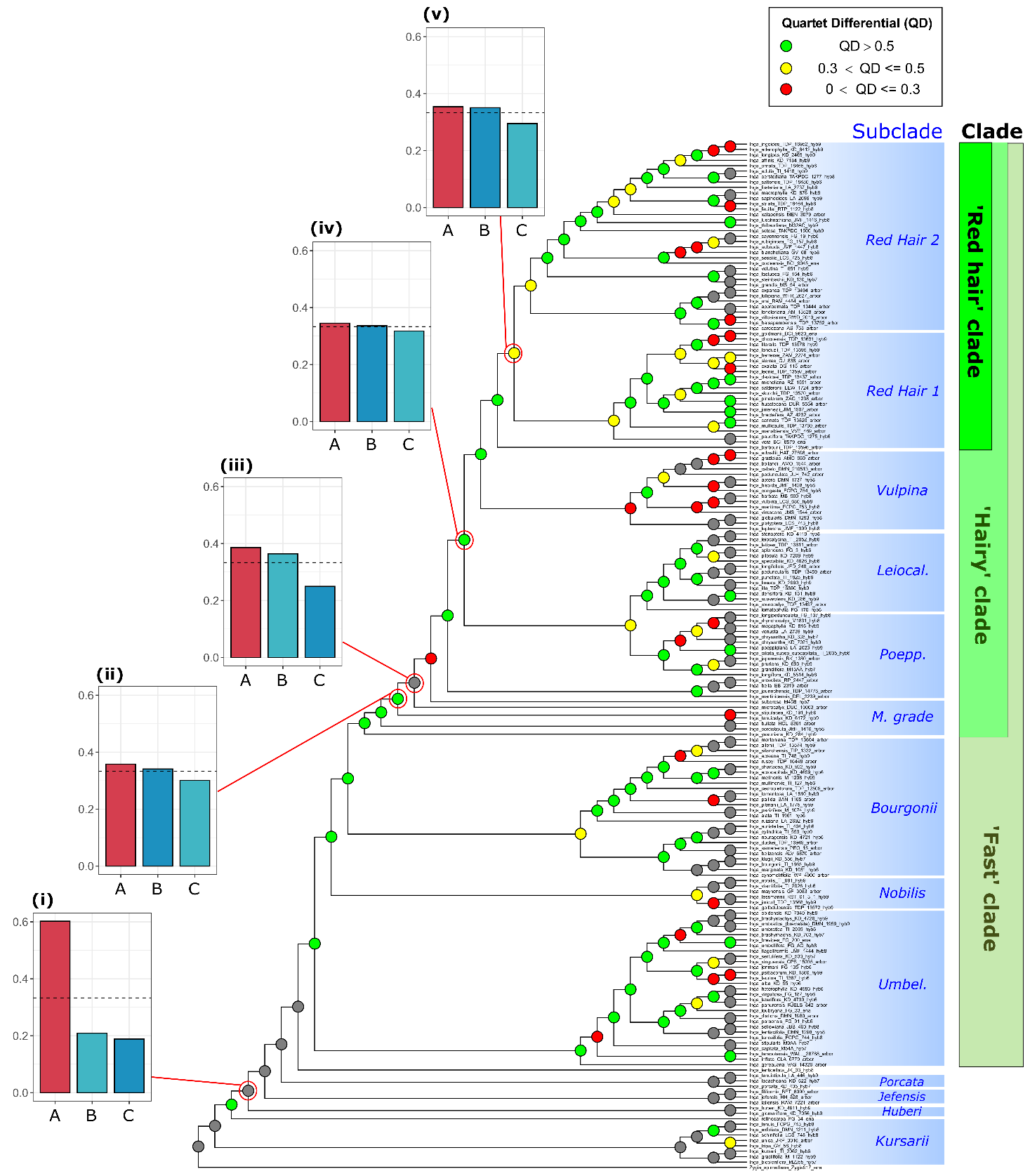

**Figure S6c:** ASTRAL tree of *Inga* ‘PPD’ dataset with paralogs removed. QD values are plotted on each node, indicating the differential between the frequency of the main topology with conflicting topologies (low QD = one incongruent topology favoured). Nodes of interest are annotated with DiscoVista plots, showing the relative proportions of different discordant topologies. Quartet frequencies are represented as bar graphs, with red bars (left) representing the main topology from the ASTRAL analysis, and with blue and turquoise bars (middle and right) representing alternative topologies. Dashed horizontal lines mark the expectation for equal frequencies of the three possible topologies (Y = 0.333), i.e. maximal gene tree conflict. Node i indicates low proportions of both conflicting alternative topologies. Nodes ii-v indicate one major conflicting topology. Subclades and generic-level clades are labelled as in Fig. 1a.

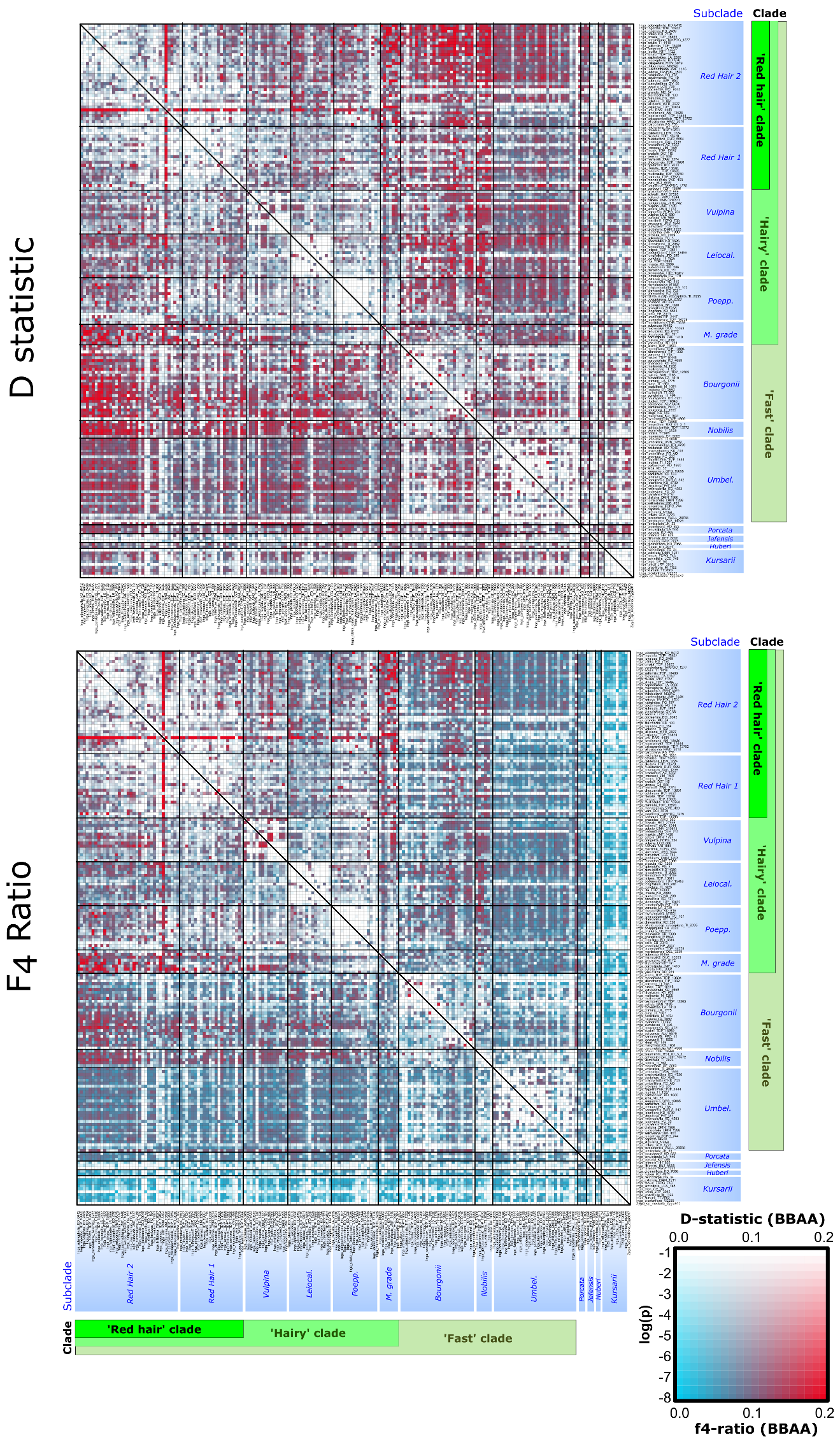

**Figure S7**: Heatmap of minimum F4-ratios plotted for the *Inga* ‘PPD’ dataset with putative paralogs removed. Taxa P2 and P3 are displayed on the *x-* and *y*-axes in the same order as in Figure 1a. The colour of each square signifies the *D-*statistic or F4 ratio estimate (blue = low estimate; red = high estimate). The saturation of these colours represents the P-value for that test (see see log(P) in inset box, bottom right). Subclades and generic-level clades are labelled as in Fig. 1a.

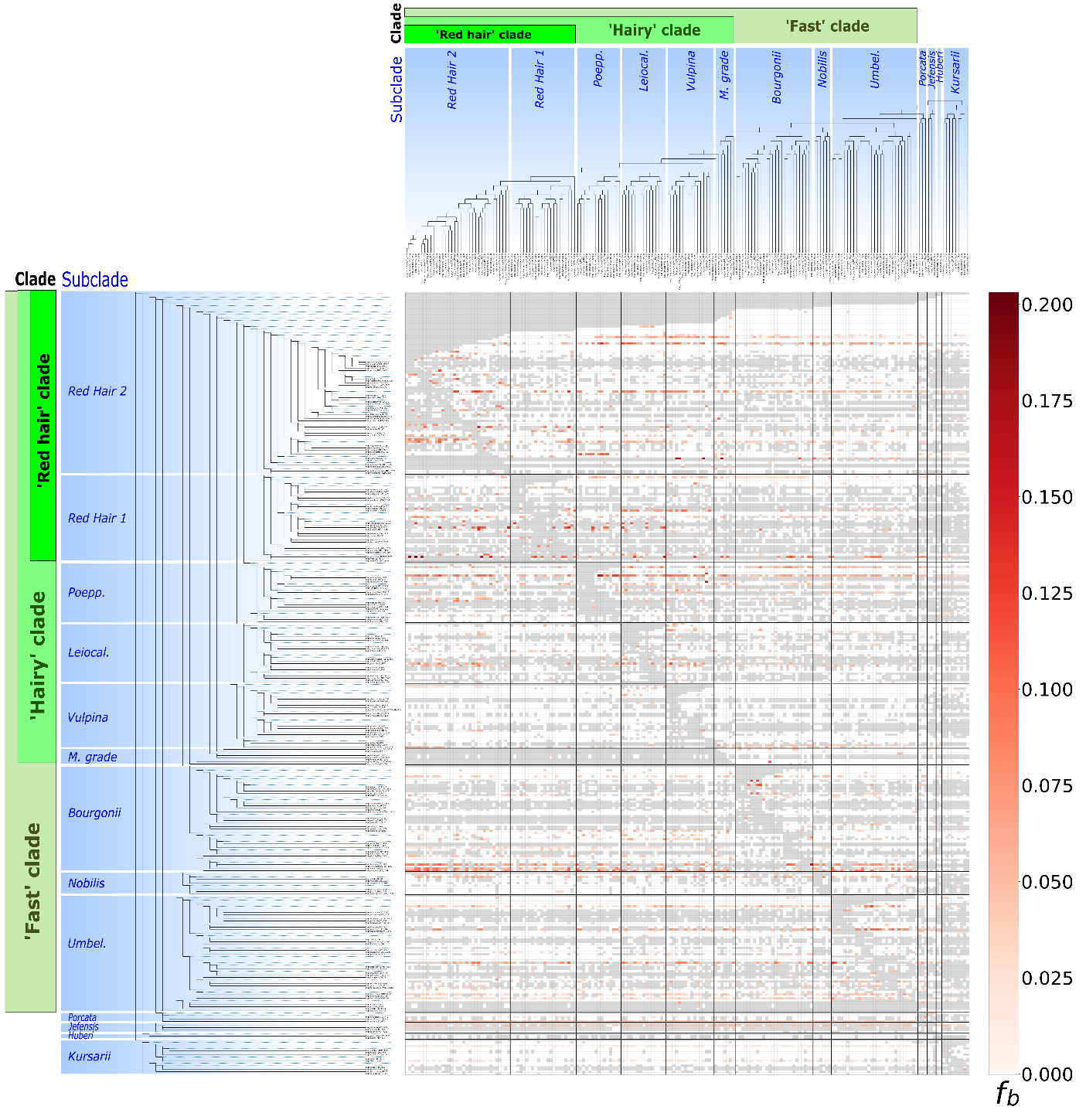

**Figure S8a**: Heatmap of ‘Fbranch’ scores plotted for the *Inga* ‘Singlesp’ dataset, representing excess allele sharing between sets of taxa and internal branches. The tree is displayed in an ‘expanded’ form along the y axis, so that all branches (including internal ones, marked as dotted lines), correspond to a row in the heatmap. The phylogenetic tree from Figure 1a is represented on the x axis in the same order as in Figure 1a such that each tip corresponds to a column in the heatmap. The colour of each square in the heat map signifies the amount of excess allele sharing (f_b_; dark red = high estimate). Subclades and generic-level clades are labelled as in Fig. 1a.

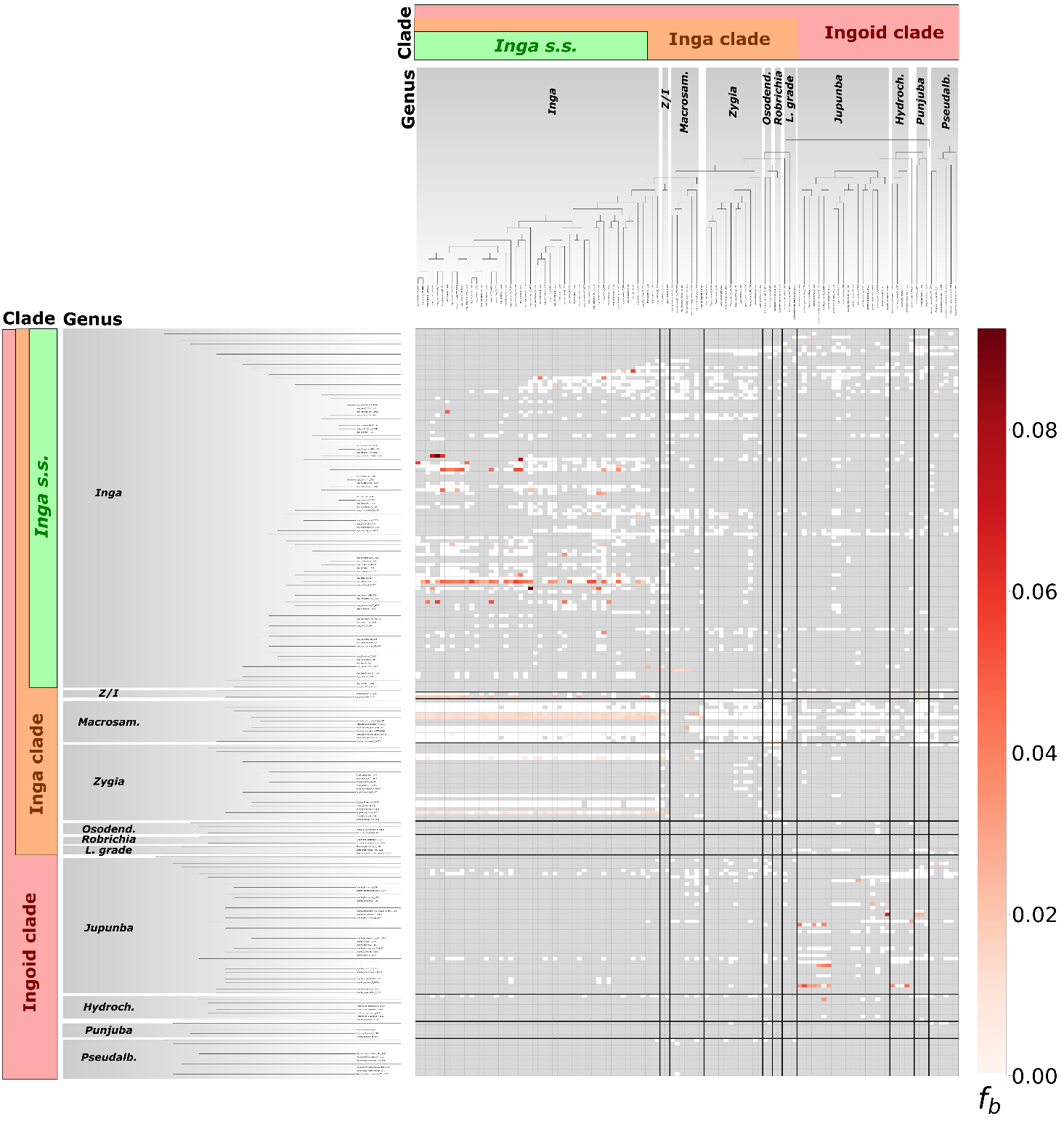

**Figure S8b**: Heatmap of ‘Fbranch’ scores plotted for the ‘Outgroup’ dataset, representing excess allele sharing between sets of taxa and internal branches, showing only pairs of taxa with significant genomic clustering of ABBA patterns to account for substitution rate variation across taxa. The tree is displayed in an ‘expanded’ form along the y axis, so that all branches (including internal ones, marked as dotted lines), correspond to a row in the heatmap. The phylogenetic tree from Figure 1b is represented on the x axis in the same order as in Figure 1b such that each tip corresponds to a column in the heatmap. The colour of each square in the heat map signifies the amount of excess allele sharing (f_b_; dark red = high estimate). Clades are annotated by genus, and then by the broader phylogenetic clades in which they are nested (Inga clade, Ingoid clade) as in Fig. 1b. In shortened genus annotations, Z/I = *Zygia/Inga*, *Macrosam*. = *Macrosamanea*, Osodend. = *Osodendron*, L. grade = *Leucochloron* grade, Hydroch. = *Hydrochorea*. Pseudoalb. = *Pseudoalbizia.*

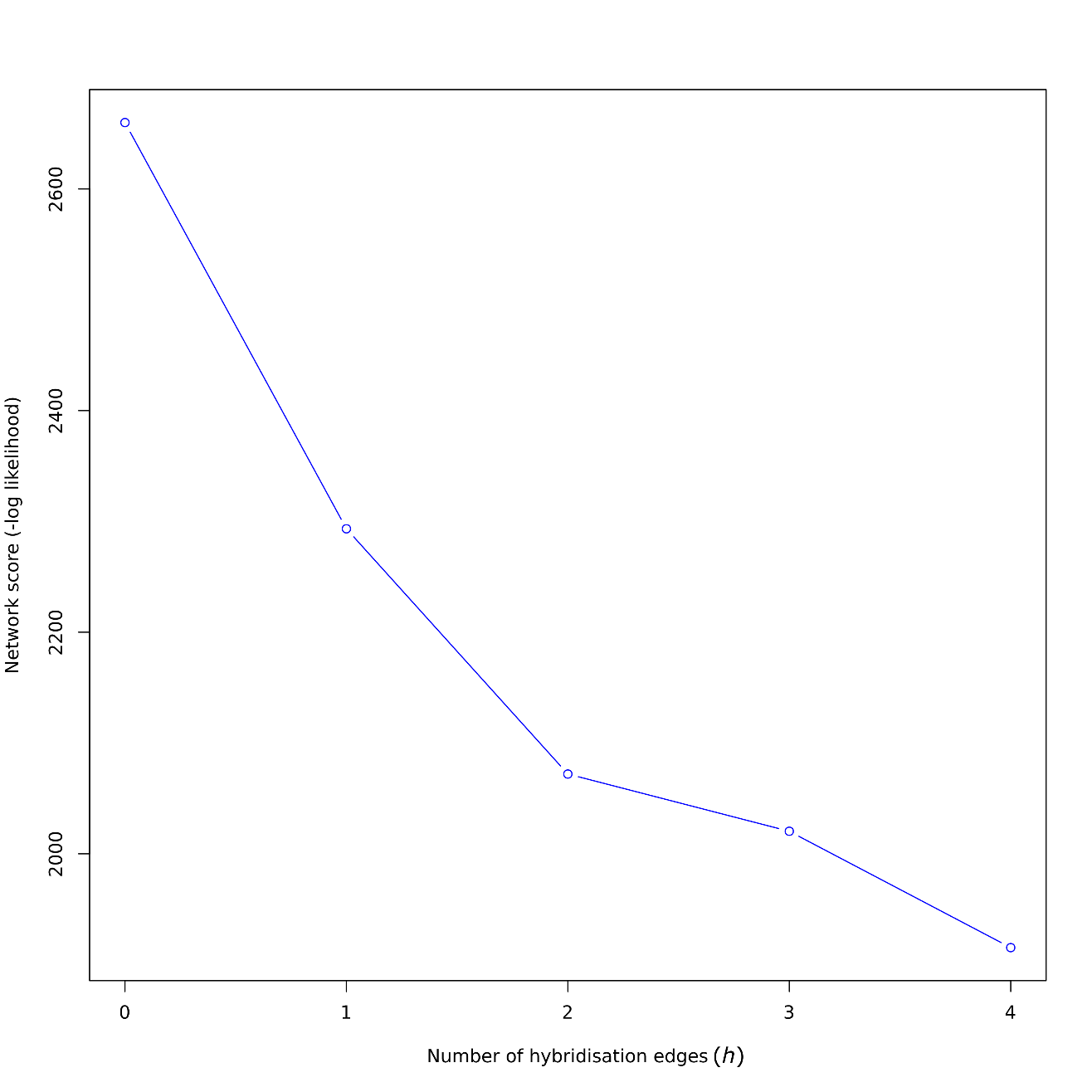

**Figure S9ai:** Negative log pseudolikelihood profile for between 0-4 hybridization events (*h)* inferred onto the *Inga* single-accession-per-species ASTRAL tree using *SNaQ!* as implemented in *PhyloNetworks*. The best-fitting number of hybridization events *(h)* is displayed as the value at which the rate of change in -log pseudolikelihood plateaus, which in this case was *h*=4.

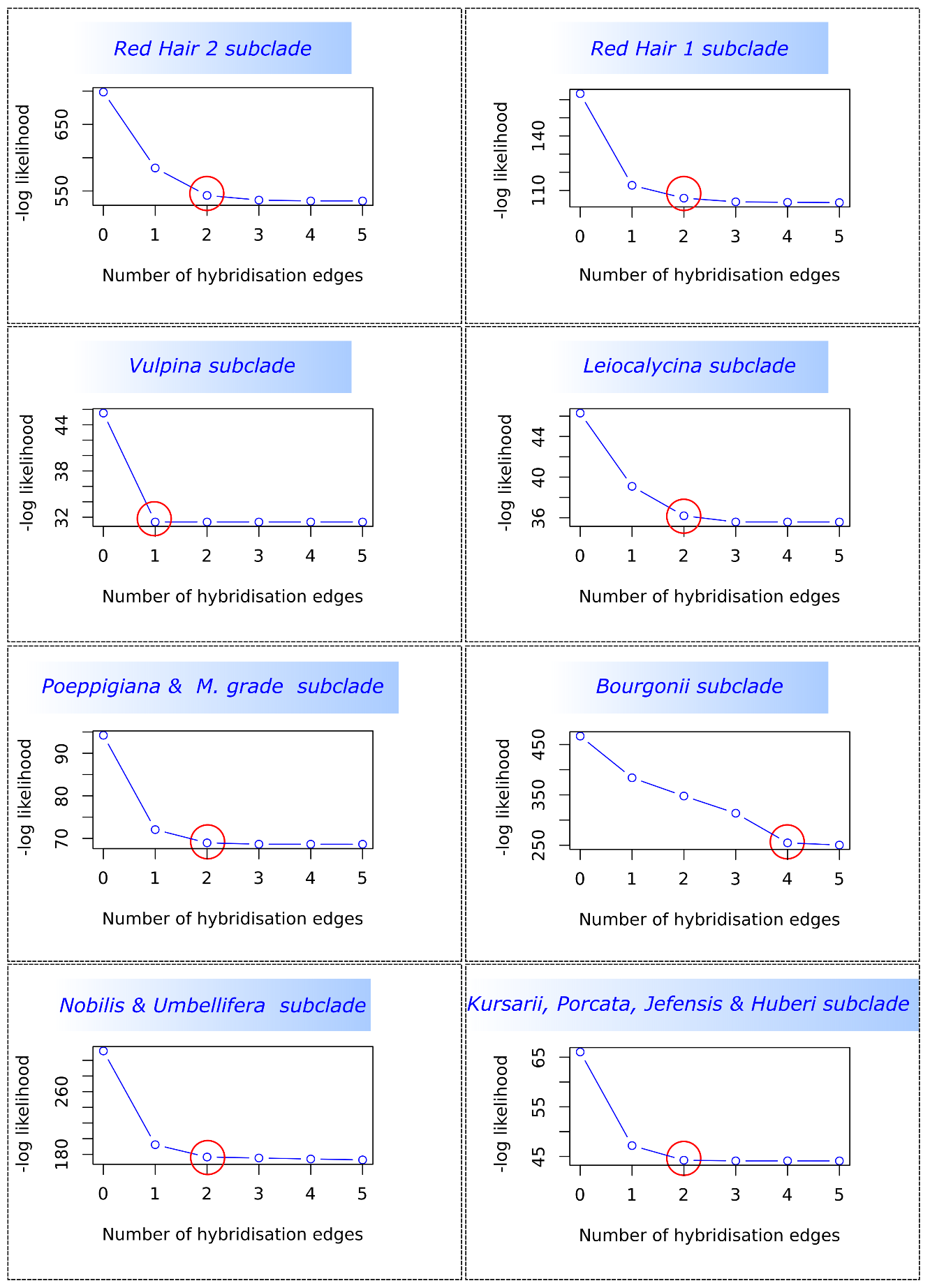

**Figure S9aii:** Negative log pseudolikelihood profile for between 0-5 hybridization events (*h)* inferred onto subclade-level trees, trimmed from the *Inga* single-accession-per-species ASTRAL tree. Networks were inferred using *SNaQ!* as implemented in *PhyloNetworks* per *Inga* subclade. The best-fitting number of hybridization events *(h)* is displayed as the value at which the rate of change in -log pseudolikelihood plateaus. The best values are circled in red for each subclade run.

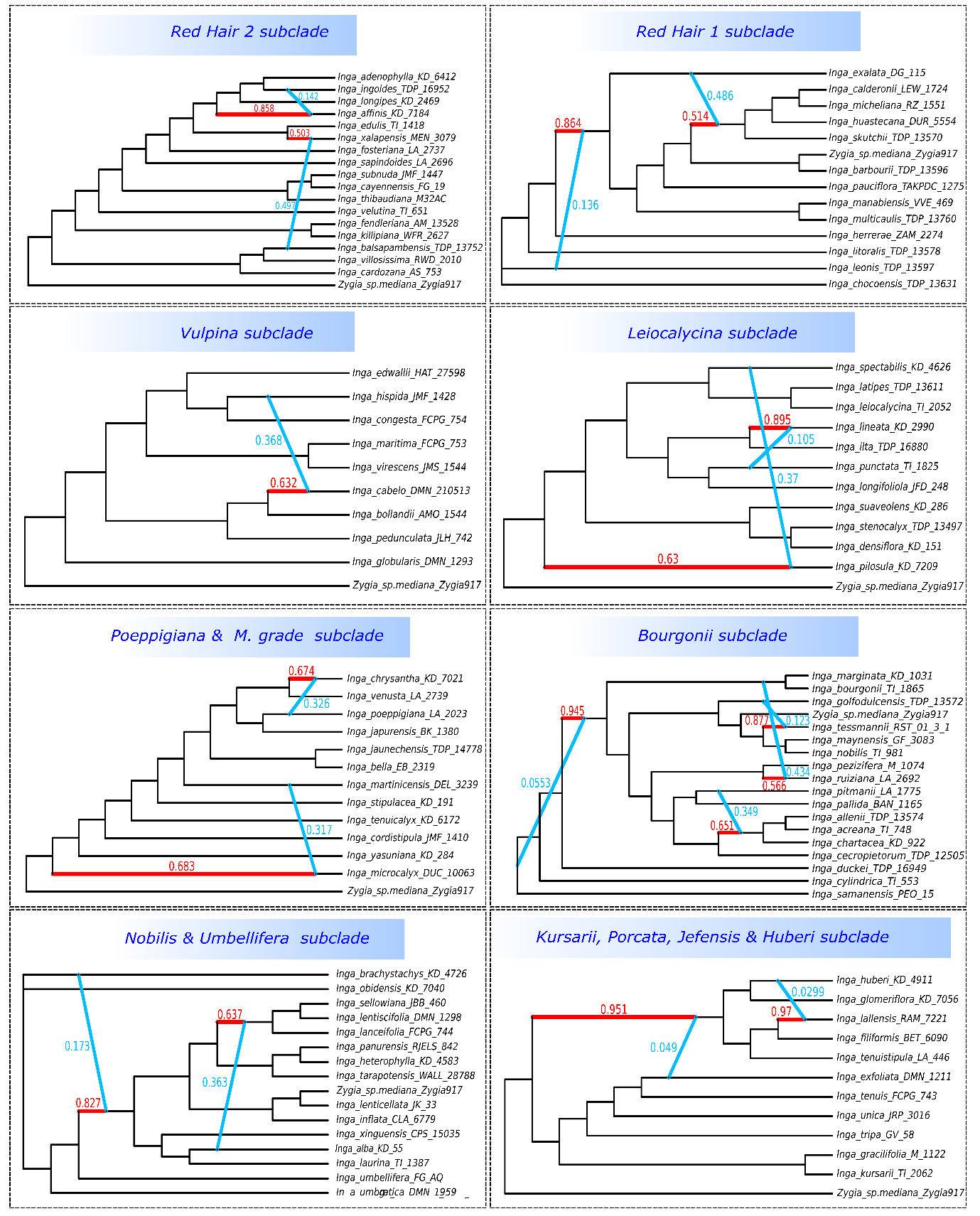

**Figure S9aii:** Networks, inferred per *Inga* subclade, using the best number of hybridisation events (*h)* indicated for each subclade in Fig. S9aii. Networks were inferred using *SNaQ!* as implemented in *PhyloNetworks* per *Inga* subclade. All networks were rooted with *Zygia* sp. ‘mediana’, but networks in which this species is not plotted bottom-most are those in which a hybridisation even involved the sister branch to the outgroup.

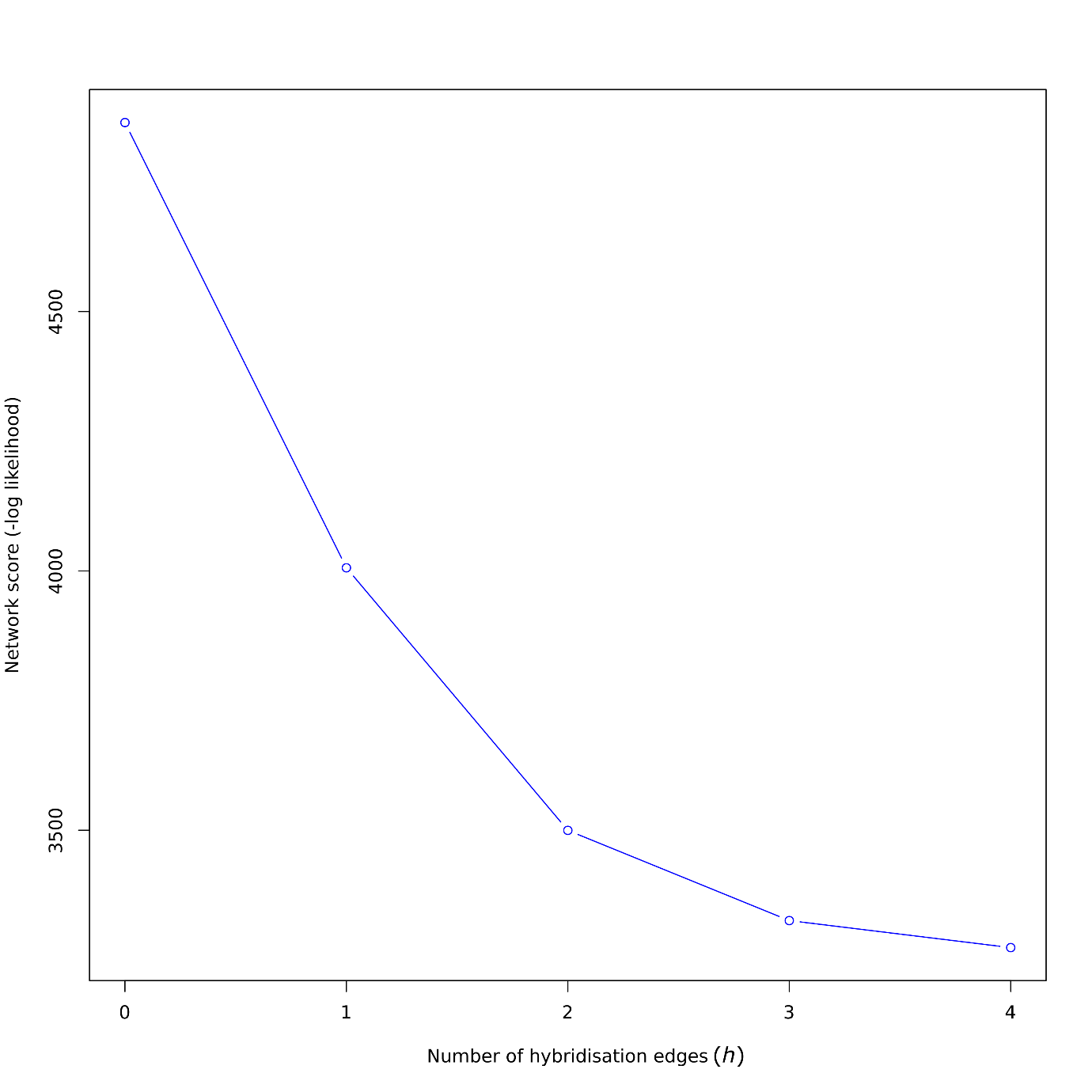

**Figure S9b:** Negative log pseudolikelihood profile for between 0-4 hybridization events (*h)* inferred onto the Outgroup ASTRAL tree using *SNaQ!* as implemented in *PhyloNetworks*. The best-fitting number of hybridization events *(h)* is displayed as the value at which the rate of change in -log pseudolikelihood plateaus, which in this case was *h*=3.

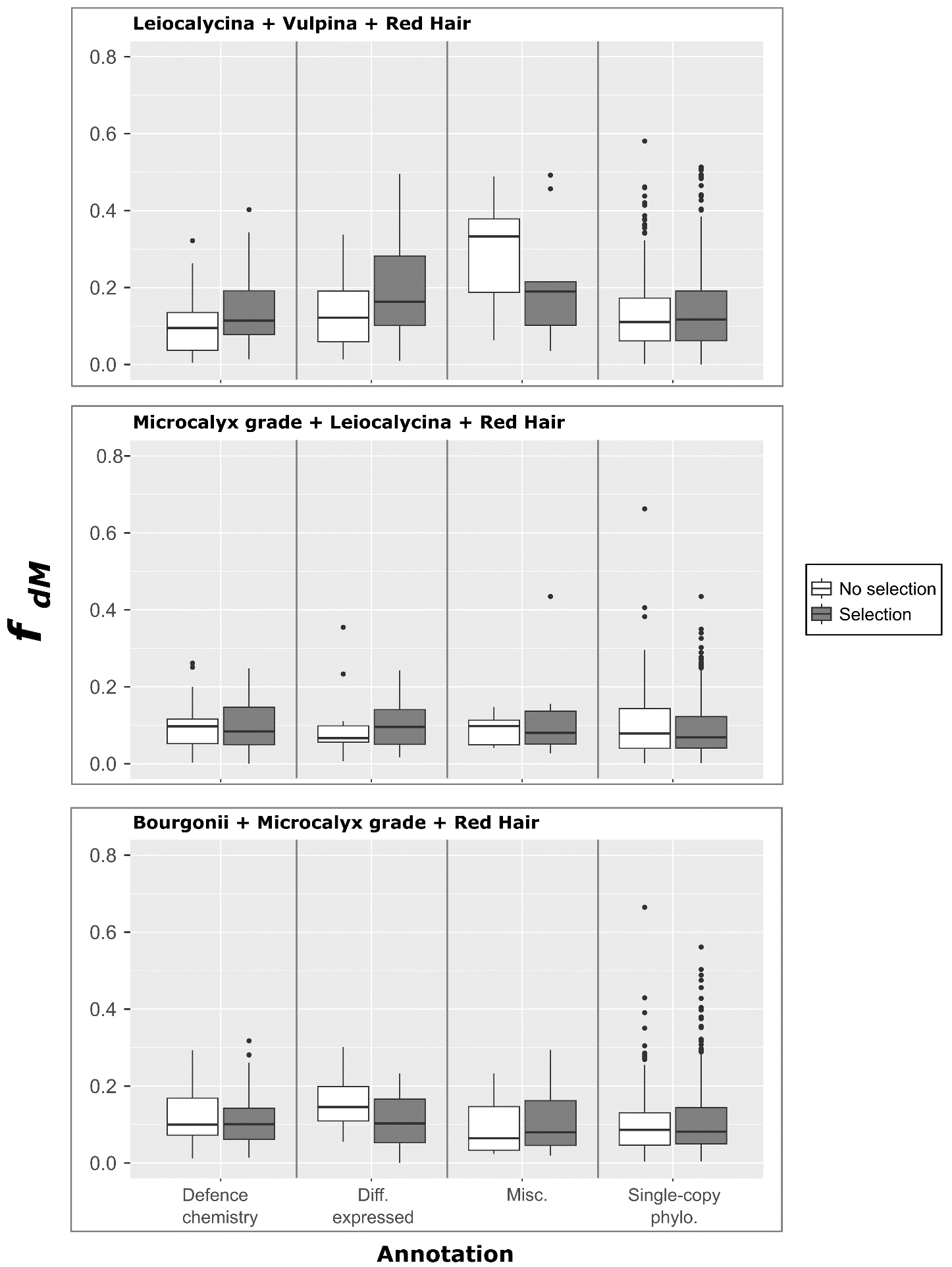

**Figure S10:** Box plots of per-locus proportion of introgression (*f _dM_*), grouped by selection result (‘under selection’ if the P-value of BUSTED analysis <0.05) and locus annotation (x-axis) for all three subclade subsets on which *f _dM_* and BUSTED analysis were carried out.

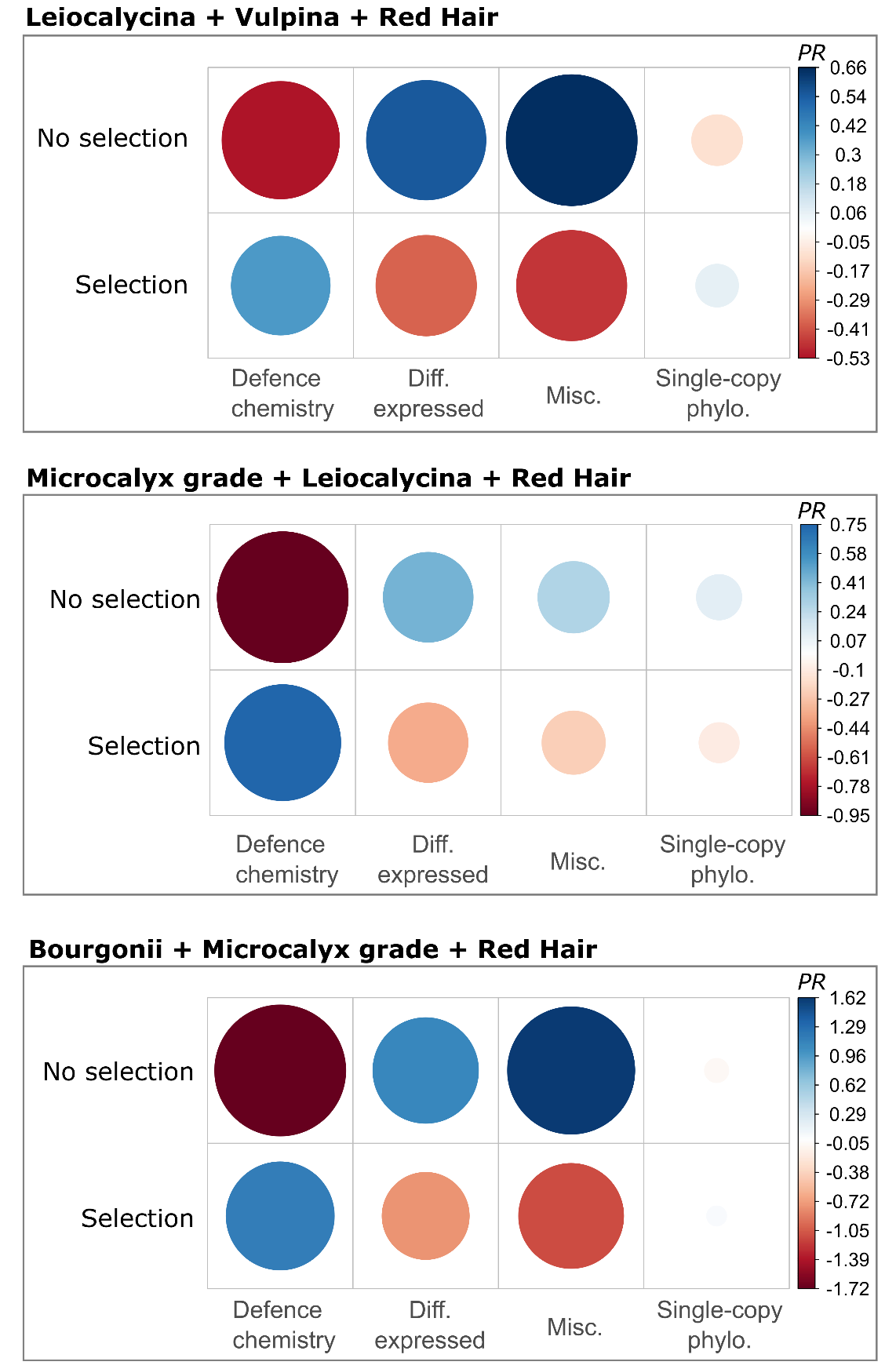

**Figure S11:** Per-locus selection (BUSTED) results summarised for all taxa and loci (title highlighted in red), as well as the three data subsets for which BUSTED and *f_dM_* analyses were performed. The correlation plots show associations between selection (‘under selection’ if the FDR-corrected P-value of BUSTED analysis <0.05) and locus annotation (Defence chemistry; Differentially expressed; Miscellaneous; Single-copy phylogenetically informative). The size of each circle represents the Pearson’s residual from χ2 analysis (*PR* in the figure legend) and hence the magnitude of the association. The colour of each circle indicates whether a proportion is above (positive = blue) or below (negative = red) the null expectation of χ2.
