## Supplementary Methods for "Rampant Reticulation in a Rapid Radiation of Tropical Trees - Insights from *Inga* (Fabaceae)"

Running the PPD pipeline to remove paralogs

We used the putative paralog detection pipeline (PPD) to identify target capture loci that were potentially paralogous within our ‘Singlesp’ *Inga* dataset. This pipeline filters sequences with a high proportion (>0.05%) of heterozygous sites, trims sites missing in >50% samples, removes hypervariable (‘noisy’) sites along a sliding window. PPD identifies paralogues by assessing whether a locus contains shared heterozygosity across >50% of taxa in the alignment. We ran the pipeline with its default settings, but changed the number of polymorphic sites allowed per sliding window to include more sequence variation (-mi 20 and -mo 40), as running the program with the default settings retained too little phylogenetic information in preliminary analyses. The resulting dataset, with putative paralogs removed, is referred to as the ‘PPD’ dataset, and was used to ascertain whether paralogous loci influenced our results.

SNP calling

We generated an input VCF file for Dtrios for each of the three datasets by calling SNPs de-novo as in the ‘Speciation Genomics’ github (<https://speciationgenomics.github.io>), using our target capture baits as the reference. First we used BWA 0.7.17 to align reads to the reference with the -M flag to mark low-quality alignments (i.e. those split across long distances) as secondary. We then sorted aligned reads with SAMTOOLS 1.13 (Danecek et al. 2021) using the ‘sort’ function. PCR duplicates were removed with picard (<https://broadinstitute.github.io/picard/>) using the MarkDuplicates function and the following settings: ‘REMOVE_DUPLICATES=true’, ‘ASSUME_SORTED=true’, ‘VALIDATION_STRINGENCY=SILENT’,’MAX_FILE_HANDLES_FOR_READ_ENDS_MAP=1000’. Finally, we called SNPs using the BCFTOOLS 1.13 (Li 2011) mpileup function with the ‘-a’ flag to add annotations for allelic depth (AD), genotype depth (DP) and strand bias (SP) to the output VCF.

D and F statistics for inferring introgression

The D statistic, also known as the ABBA/BABA test, uses asymmetry in gene tree topologies to quantify introgression between either of two lineages which share a common ancestor (P1 and P2) and one other lineage (P3) that diverged from the common ancestor of P1 and P2 (Green et al., 2010; Durand et al., 2011). This is done by comparing patterns of derived alleles and alleles from an outgroup (O) across sets of four-taxon gene trees. The D statistic is also known as the ABBA/BABA test because, assuming the outgroup allele is labelled ‘A’ and the derived copy is labelled ‘B’, these are the patterns of alleles present in the four-taxon trees with the order [P1, P2, P3,O]. If the proportion of these alleles is significantly different from 1:1 (as would be expected under ILS), then hybridization has occurred between either [P1 or P2] and [P3]. If D < 0, then P1 and P3 have a history of introgression, and if D > 0, then P2 and P3 have a history of introgression. For more information, see equation 2 of the Dsuite publication (Malinsky et al. 2021).

The F4-ratio is calculated in a similar fashion to the D-statistic, based on four populations, here with the topology topology (A,B),(C,D). However, the F4 ratio differs from the D-statistic in that it also assumes the existence of an extra admixed population, in between (A,B) and (C,D), testing for departure from expected allele frequencies between populations assuming the null hypothesis (of no introgression) is true. Specifically, the F4 ratio can estimate the proportion of introgressed genetic variation (i.e., F4 ratio values >0) by calculating the product of the difference of allele frequencies between A and B, and between C and D, divided by the product of the difference between A and C and between C and D (equation 5, Malinsky et al. 2021). As an extension of this, to account for correlated F4-ratio scores of related branches where descendent species inherit introgressed variation, the Fbranch metric calculates a median of F4 ratio scores across related branches (equation 6, Malinsky et al. 2021).

Finally, to estimate per-window proportions of introgressed variation (i.e. excess allele sharing), the fd metric assumes three populations and an outgroup with the relationship (((P1, P2), P3), O) and essentially involves subtracting the frequency of excess of derived alleles in either P2 or P1 (i.e., subtracting the number of ABBA from the number of BABA patterns) across the genomes of four populations (the numerator), and then normalising it by dividing this value by the total number of sites (the demoninator). FdM, which was derived from fd, goes one step further by dividing the numerator by either by one of two denominators: if P2 has more derived alleles, then fdM=fd=S(P1;P2;P3;O)/S(P1,PD,PD,O), but if P1 has more derived alleles then fdM=S(P1;P2;P3;O)/-S(PD,P2,PD,O). This means that the fdM statistic is centred around 0, with the sign of the score (+/-) detailing which population pair experience introgression – P1/P3 or P2/P3 (see Equation 8, Malinsky et al. (2021).

PhyloNetworks accession selection

For the Singlesp and PPD datasets, we down-sampled the same 27 *Inga* species. These species were chosen to represent subclades within *Inga*, with representative species chosen based on those with the highest recovery success within a subclade (recovery success per-accession is summarised in Supplementary Information Table S3, available on Dryad). We also included two species with strong evidence of introgression in other analyses (*I. microcalyx* and *I. barbourii*). We applied the same methods of selecting species with the best locus recovery from each subclade ‘per-subclade’ PhyloNetworks analyses.

For the down-sampled ‘Outgroup’ dataset, we mainly selected species within the *Inga* clade since we were most interested in contextualising introgression in *Inga* by examining reticulation events subtending it. Once again, we used proportional per-clade sampling of species based on those with the highest recovery success.

Subclade selection for inferring per-locus selection and introgression

We assessed whether chemical defence loci experienced elevated introgression (*f_dM_* in Dsuite) and selection (BUSTED) relative to other loci in our target capture dataset. We performed the analysis on three subsets of our ‘Singlesp’ dataset, containing three different combinations of taxa to estimate the amount of per-locus introgression between the subclades with the most evidence of introgression in our Dsuite and PhyloNetworks analyses.

We had to use these three data subsets because estimating per-locus *f _dM_* for all loci across all combinations of *Inga* species was computationally impractical. We aimed to assess whether per-locus introgression (and selection) was most prominent in defence chemistry loci at the base of *Inga* subclades, under the hypothesis that these introgression events may have transferred adaptive loci between species in these subclades.
